# Regenie.QRS: computationally efficient whole-genome quantile regression at biobank scale

**DOI:** 10.1101/2025.05.02.651730

**Authors:** Fan Wang, Chen Wang, Tianying Wang, Marco Masala, Edoardo Fiorillo, Marcella Devoto, Francesco Cucca, Iuliana Ionita-Laza

**Affiliations:** Department of Biostatistics, Columbia University, New York, US; Department of Statistics, Colorado State University, Fort Collins, US; Institute for Genetic and Biomedical Research, National Research Council, Italy; Department of Biomedical Sciences, University of Sassari, Sassari, Italy; Department of Statistics, Lund University, Lund, Sweden

## Abstract

Genotype-phenotype associations can be context-dependent and dynamic in nature leading to heterogeneity of genetic effects across different parts of the phenotype distribution. Quantile regression, an alternative to linear regression for continuous phenotypes, is particularly well suited for detecting and characterizing heterogeneous genotype-phenotype associations. Here we propose a novel and computationally efficient whole-genome quantile regression technique, Regenie.QRS, for biobank-scale GWAS data with genetic structure. Our approach first estimates the polygenic effect, and then incorporates this effect as an offset in the non-mixed quantile regression model. Our simulations demonstrate robust control of type I error and higher power to detect heterogeneous associations relative to linear regression in GWAS, and improved power over the marginal quantile regression tests. We present applications using data from the UK Biobank and the ProgeNIA/SardiNIA project, where we show the advantages of Regenie.QRS in identifying and characterizing heterogeneous genetic effects. To cite just one interesting example, using quantile regression we are able to show that even though variants at the *G6PC2* locus increase glucose levels, their effects are much stronger at lower quantiles of glucose level distribution than at higher quantiles, showing that *G6PC2* serves as a guardian against low glucose levels without driving dangerous hyperglycemia, which may explain the lack of association with diabetes risk.

## Introduction

Genome-wide association studies (GWAS) have been the dominating approach to identify genetic variants associated with complex phenotypes. For many traits, such studies have started to provide important knowledge about genetic architecture, disease pathways and numerous genetic loci for follow-up. Despite these significant advances, we have limited understanding of how genotype-phenotype associations vary across contexts, hindering our ability to move closer to a more mechanistic understanding of these associations. Disease-related variants have a tendency to be context-dependent, e.g. causal variants for kidney function have been shown to preponderantly fall into cis regulatory elements that are active in a few kidney cell types including kidney tubule epithelial and podocyte^1^. The challenging aspect of modeling context-dependency is that oftentimes the right context is not known ahead of time and it is difficult to collect comprehensive data across multiple environments or contexts.

The conventional GWAS approaches are based on linear regression models (LR). These models have been widely adopted for two primary reasons: (1) they are easy to implement and interpret, and (2) GWAS are typically based on meta-analyses of multiple individual studies, and hence simplified models are practical for implementation and meta-analysis across studies. However, LR methods assume homogeneity of effects across contexts and therefore offer no insight into how the resulting associations vary across contexts, and can even miss these context-dependent associations. Quantile regression (QR) is a statistical technique that is well suited to identify and characterize such context-dependent associations without directly modeling interactions with context. The key idea is that unmodelled context-dependency can lead to heterogeneity in genetic effects across different quantiles of the phenotype distribution, and therefore the effect may be stronger at some quantiles relative to other quantiles. QR works by fitting a series of linear regression models at various quantile levels of the phenotype, allowing for the estimation of genetic effects across the entire trait distribution.

QR has wide applications across diverse fields such as economics and finance, medicine, epidemiology, social sciences, etc. Its applications to genetics have been limited due to a relative lack of familiarity with QR among geneticists and a lack of scalable QR techniques that can handle complexities of large scale GWAS data, such as the (ultra) high-dimensional data in biobanks and complex genetic structure with correlated individuals. Here we propose to develop scalable QR methods for biobank scale data with related individuals. Although QR is only applicable to quantitative traits, such traits including disease biomarkers and intermediate cellular phenotypes are being collected at increasing scale due to their importance in moving us closer to causal mechanisms in genetic association studies and in phenotype prediction. Similarly, for plants and animals quantitative traits are highly relevant as they often represent key phenotypes that influence productivity, survival and adaptation.

In recent work we have applied QR to gene expression phenotypes in GTEx and various endophenotypes in UK Biobank (UKBB)^2;3^ under the assumption of independence among samples. Although there are related individuals in UKBB, we implemented a simple approach whereby we removed related individuals from the study. However, removing individuals to ensure independence is not ideal, as it may lead to loss of power and increased false positive rates due to possible residual genetic structure in the data. Quantile mixed effect models provide a natural solution to conduct quantile regression with related samples, and a number of such models have been proposed in the context of longitudinal or clustered data. These include distribution-free approaches ^4;5;6;7^, and likelihood-based approaches based on the asymmetric Laplace density (ALD) ^8;9;10^. The likelihood-based approach is motivated by the fact that minimizing the pinball quantile loss function in QR is equivalent to maximizing the likelihood under the Laplace distribution function. Among the ALD-based models, Geraci et al. ^9^ proposed a Monte Carlo EM procedure for subject-specific random intercepts to avoid the evaluation of a multidimensional integral, which is computationally burdensome. Subsequently, Geraci et al.^11;12^ explored alternative computational techniques based on Gaussian quadrature and non-smooth optimization algorithms. More recently, Galarza et al. ^13^ adopted a stochastic approximation of the EM algorithm. Despite these developments, these methods suffer from scalability issues when applied to large-scale GWAS with related samples.

Here we propose to develop a computationally efficient framework to estimate polygenic effects, and then incorporate these effects as offset in the non-mixed quantile regression model. In GWAS, the random effect represents the polygenic effect captured through sample relatedness, often represented as SNP-derived genetic relationship matrix (GRM). Specifically, we propose a novel quantile rank score (QRS) test, Regenie.QRS, based on the machine learning technique of stacked regression, similar to the one implemented in the whole-genome linear regression based method Regenie ^14^. Regenie was shown to be a powerful GWAS method while also being computationally efficient. A more traditional approach to estimate the polygenic effect is via best linear unbiased prediction (BLUP) and is commonly used in animal and plant studies. While BLUP is powerful, it can be computationally intensive. This computational burden can be mitigated by using iterative solvers such as the preconditioned conjugate gradient (PCG) descent^15^, which avoids direct matrix inversion and enables scalability to large datasets. For the sake of comprehensiveness, we also discuss a BLUP based approach and compare it against Regenie.QRS. We further derive the asymptotic linear representation and consistency of quantile regression estimation based on polygenic predictions, demonstrating that the theoretical requirements for accurate polygenic predictions are met by various estimators, including Regenie and BLUP.

Through simulations, we demonstrate that Regenie.QRS has correct type-1 error and is more powerful than Regenie when effects are heterogeneous across quantile levels. Furthermore, Regenie.QRS is more powerful than the regular marginal QRS test, while controlling type-1 error under complex genetic structure. We apply Regenie.QRS to 28 quantitative traits using data from UKBB and a second independent dataset from the ProgeNIA/SardiNIA study, a longitudinal cohort of individuals from Sardinia. Results demonstrate that our framework effectively uncovers genetic associations with heterogeneous effects across different quantile levels, providing more nuanced insights into complex genotype-phenotype associations that are not possible in the conventional linear framework.

## Methods

### Linear quantile mixed model

Let **Y** denote the *N ×* 1 phenotype vector, and **G** represent the *N × M* standardized genotype matrix. **G**_**j**_ represents the *N ×* 1 genotype vector for SNP *j*. Let **Z** represent the *N × C* matrix of covariates (including an intercept). In the classical linear mixed GWAS model ^14;16;17^,

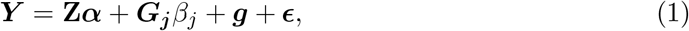

where **g** is the polygenic effect accounting for relatedness and population structure with 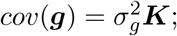 ***K*** is the GRM, and 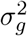 is the phenotypic variance attributed to the causal SNPs. ***ϵ*** is the vector of residuals 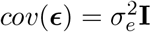. Although the computation of ***K*** should be based on the true causal variants, in practice we estimate the GRM by selecting a subset of variants^16;17;18^.

To allow genetic effects and effects from other covariates to depend on the quantile *τ ∈* (0, 1) of interest, we consider the following model for the conditional quantile functions of the phenotype *Y*_*i*_ for the *i*-th individual:

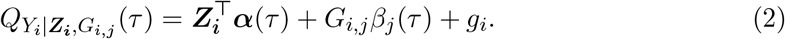

We assume here that *g*_*i*_ has a pure location shift effect on the conditional quantiles of the phenotype, and that its covariance (which captures relatedness among individuals) does not depend on the quantile.

To fit model (2), we adopt a two-step procedure. First, we estimate the polygenic effects to obtain *ĝ*, and then we perform estimation or testing of *β*_*j*_(*τ* ) after substracting the estimated polygenic effects. To estimate the polygenic effects, we consider two approaches: one based on the recent machine learning method Regenie ^14^, and one based on the traditional statistical approach BLUP. To avoid LD contamination, we adopt the Leave-One-Chromosome-Out (LOCO) scheme^19^ so that the polygenic prediction is calculated from SNPs on all chromosomes except the one containing the tested SNP. We provide the details of the two methods for polygenic effect estimation in the next section.

The estimation of *β*_*j*_(*τ* ) for SNP *j* is obtained by minimizing the following quantile loss function:

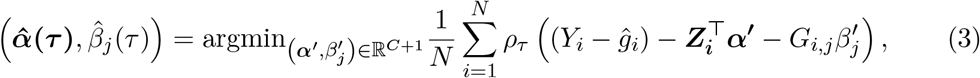

where *ρ*_*τ*_ (*µ*) = *µ*(*τ − I*(*µ* < 0)) is the pinball loss function ^20^. In Supplemental Material (section on *Asymptotic properties*) we demonstrate that, under mild conditions, the quantile regression estimation in (3) based on polygenic predictions *ĝ* is consistent.

For hypothesis testing, a commonly used tool is the QRS test^21^. We define ***Y***_***resid***_ = ***Y*** − ***ĝ***. The QRS test to test SNP *j* for a fixed quantile level *τ* is based on

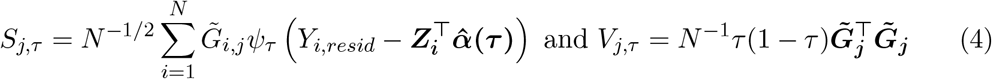

where 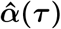 is obtained by minimizing the function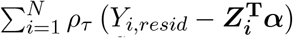, and 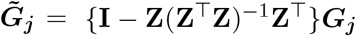.Under the null hypothesis 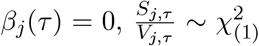. It is important to note that the asymptotic distribution of the test statistic is independent of the specific distribution of the phenotype. Therefore, this approach can be applied to any phenotype without the need for a pre-transformation to achieve normality. For each SNP, we conduct QRS across equally spaced quantile levels and combine quantile specific p-values via Cauchy combination ^22^.

### Polygenic predictions using Regenie

Here we provide an overview of Regenie, an approach inspired by stacked regressions in machine learning ^23^, which can predict polygenic effects. Regenie assumes the following whole-genome regression model with *M* SNPs:

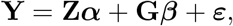

where 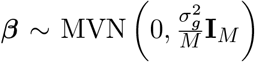 and 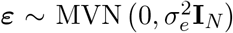. This is equivalent to the standard infinitesimal model:

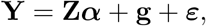

with 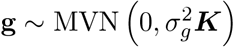, where ***K*** = **GG**^*T*^ */M* is the GRM.

Let 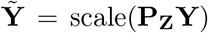 and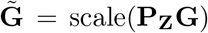, where **P**_**Z**_ = **I**_**N**_ − **Z(Z**^*T*^**Z**)^−1^ **Z**^*T*^ , and scale(*·*) denotes standardization to have unit variance. That is, 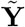 and 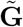 represent the phenotype and genotype residuals after regressing out covariate effects and scaling to have unit variance. First, we partition the genome into *B* consecutive blocks. Given a genotype block 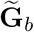 of *M*_*b*_ SNPs, we fit ridge regression for the following model

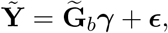

where ***ϵ*** = (*ϵ*_1_, … , *ϵ*_*N*_ )^*T*^ with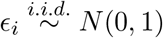. The ridge regression estimate for the genetic effects ***γ*** with ridge parameter *λ* is given by

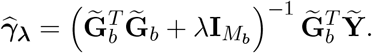

Assuming a Bayesian prior where SNP effect sizes follow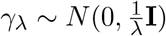, the total genetic variance is 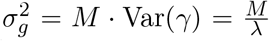. Setting this equal to SNP heritability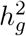, we solve for 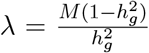. We select *R* evenly spaced values of 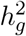 within the interval [0.01, 0.99]. For each block of size *M*_*b*_, we end up with a much smaller set of *R* predictors,

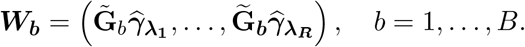

Let ***W*** = (***W***_**1**_, … , ***W***_***B***_) be the resulting *N × BR* matrix obtained from fitting the ridge regressions on all the *B* genotype blocks. We then combine the predictors in ***W*** . Specifically, we re-scale the predictors in ***W*** to have unit variance and apply ridge regression again, which corresponds to the following model

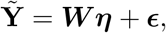

where ***ϵ*** = (*ϵ*_1_, … , *ϵ*_*N*_ )^*T*^ with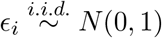. Then we obtain the following estimator

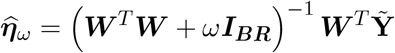

for a range of shrinkage parameters *ω* and choose a single best value using K-fold cross-validation.

The result of the model fit is 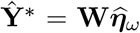. To produce polygenic predictions for each chromosome using the LOCO scheme, we compute the predictions by setting the entries in 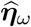 corresponding to blocks in the left out chromosome to 0. This approach is less computationally burdensome than a more exact LOCO scheme, which would involve leaving out all SNPs within the tested chromosome before partitioning the genome into blocks, and seems to work well in practice based on simulations. Regenie accommodates parallel analysis and requires only local segments of the genotype matrix to be loaded into memory.

Let *ĝ* denote the prediction from Regenie for 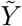 by LOCO scheme. Then we perform QRS by

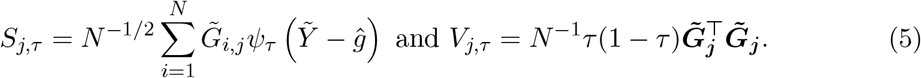

### Polygenic predictions using BLUP

The traditional BLUP for the random effect in model (1) can be written as

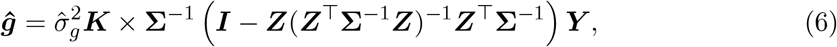

where 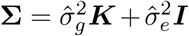 , and the variance components 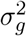 and 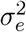 are estimated by restricted maximum likelihood (REML). As in most existing linear mixed model-based association tools ^24;25;16^, we estimate ***ĝ*** under the null model *β*_*j*_ = 0 to avoid repeatedly running the random effects estimation for each target variant. In addition, Yang et al.^19^ pointed out that including the candidate SNP while calculating the GRM ***K*** could decrease power and recommended including all SNPs except for the candidate SNP and SNPs in LD with it. This LOCO scheme has been efficiently implemented in the GCTA software, and we adopt it here as well.

The calculation in (6) involves the inversion of a large matrix **Σ**, which is computationally prohibitive in large scale GWAS. The inversion is based on Cholesky decomposition which requires 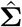 to be positive definite ^16^, or eigen-decomposition^26^. Both of these operations are *O*(*N* ^3^) for dense matrices ^27^. Although this time complexity can be greatly improved by leveraging sparsity, we adopt here an alternative estimation method based on the preconditioned conjugate gradient descent as proposed in SAIGE^28^. This method avoids direct inversion of **Σ**, and instead iteratively solves a linear system, reducing the computational cost to *O*(*Ns*), where *s* is the number of SNPs used to compute the GRM. It also stores binary genotypes instead of requiring the pre-computed GRM as an input.

We compared polygenic predictions from Regenie and BLUP in simulations and show that they are fairly strongly correlated (Fig. S1). In Supplemental Material (section on *Asymptotic properties*), we also derive the asymptotic linear representation and consistency of quantile regression estimation based on polygenic predictions, demonstrating that the theoretical requirements for accurate polygenic predictions are met by various estimators, including Regenie and BLUP.

## Results

### Simulations

We conducted simulation studies using UKBB (unimputed) genotype data. Specifically, we selected individuals of white British ancestry with available genotype data and applied quality-control filters using PLINK2 ^29;30^. These filters included: excluding individuals with more than 10% missing genotypes, retaining only SNPs with a minor allele frequency (MAF) of at least 0.05, considering only SNPs with a genotyping rate of at least 90%, and removing SNPs with a Hardy-Weinberg equilibrium p-value below 10^−4^. After applying these quality-control steps, we retained a total of 409,627 individuals and 341,257 SNPs.

We conducted simulations using 409,627 individuals and also considered extreme scenarios involving subsets of more closely related individuals. Specifically, we focused on first-degree relatives in UKBB, identified using the software KING ^31^, resulting in a dataset of 46,887 individuals with first-degree relationships. We simulated phenotypes under several common distributions including normal and heavy-tailed distributions. Additionally, we assumed that the effects of genetic variants could be non-additive and heterogeneous across different quantiles of the phenotype distribution. For hard-called genotype data (0, 1, 2), missing genotypes were set to the most common allele by default in PLINK2. We compared the following methods in terms of power and false positive rate (FPR): QRS, Regenie, Regenie.QRS, and BLUP.QRS. In what follows, all causal SNPs are randomly selected from odd chromosomes and hence FPR can be computed based on null SNPs on even chromosomes containing no causal variants. For all analyses, we include the first 10 principal components (PCs) computed by FlashPCA^32^ as covariates.

### Homogeneous setting - Gaussian linear model

We simulate phenotypes by the following model:

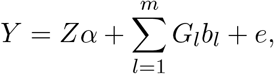

where *G*_*l*_ are genotypes for the *l*th causal variant with effect *b*_*l*_ generated from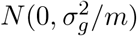. The value of 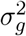 is set to be 0.6. The number of causal variants *m* is set to be 10,000. The covariate *Z* is generated from *N* (0, 1); *α* is generated from *N* (0, 0.3); *e* is a vector of residual effects, randomly generated from *N* (0, 0.1). Thus, the total phenotypic variance is set to be 1.

### Simulations under departure from Gaussian linear model

We also consider models with different error distributions and heterogeneous effects of genetic variants across quantile levels.

1. **Model with** *t*_(2)_ **errors**. We consider the homogeneous linear model above with a heavy tail error distribution *t*_(2)_. Specifically, we generate *Y* by

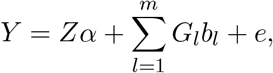

where *e ∼ t*_(2)_ distribution; *α ∼ N* (0, 1);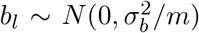. We set the value of 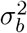 such that the heritability calculated based on the empirical variance of *e* equals 0.6.
2. **Model with dominance effects**. We next consider a model with both additive and dominance effects as follows:

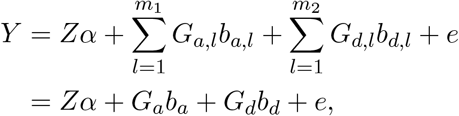

where *G*_*a*_ represents the standardized genotype matrix for *m*_1_ = 10, 000 causal SNPs corresponding to additive coding (0, 1, 2); *G*_*d*_ represents the standardized genotype matrix for *m*_2_ = 100 causal SNPs corresponding to dominance effect coding (*−p/*(1 *− p*), 1, −(1 *− p*)*/p*). The components of the vector *b*_*a*_ are independently sampled from 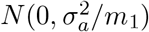,where *m*_1_ = 10,000. Similarly, the components of the vector *b*_*d*_ are independently sampled from 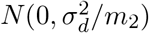, where *m*_2_ = 100. The vector of residual effects *e* is generated from *N* (0, 0.1). The covariate *Z* is generated from *N* (0, 1), and *α* is generated from *N* (0, 0.1). We set the values of 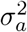 and 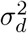 to 0.4, ensuring that the total phenotypic variance equals 1.
3. **Model with heterogeneous effects**. We consider a model with both additive effects and local effects only at upper quantiles as follows:

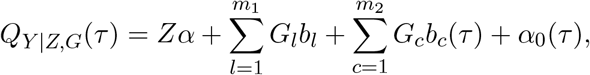

where *Z* is generated from *N* (0, 1), and *α* is sampled from *N* (0, 0.1). The values of *b*_*l*_ are generated from 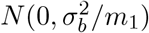 with 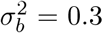 and *m*_1_ = 10,000. *G*_*c*_ represents the standardized genotype matrix for an additional *m*_2_ = 100 SNPs with local effects. When *τ >* 0.7, *b*_*c*_(*τ* ) is defined as *b*_*c*_ *× τ* , where *b*_*c*_ *∼ N* (0, 0.1*/m*_2_); otherwise, *b*_*c*_(*τ* ) = 0; *α*_0_(*τ* ) is the quantile function of the error and the error distribution 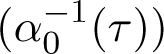 is set to be *N* (0, 0.5), such that the variance of *Y* equals 1 at quantiles below 0.7 in the absence of heterogeneous effects. To simulate *Y* , we use the inverse quantile approach.

FPR is defined as the proportion of significant SNPs among the 171,159 null SNPs located on even chromosomes, averaged over 100 replications. QRS exhibits widespread type I error inflation across multiple significance thresholds and all four genetic models (Tables 1-2, S1-S6). For example, with over 400,000 samples and at a significance threshold of 10^−6^, QRS reaches an empirical type I error rate of 5.48 *×* 10^−6^ under the homogeneous model, and 1.49 *×* 10^−5^ under the *t*_(2)_ model (Table 1). Similarly, QRS reaches an empirical type I error rate of 2.38 *×* 10^−6^ at *α* = 10^−8^ under the homogeneous model (Table S1). BLUP.QRS exhibits less inflation than QRS but still shows notable type I error inflation across all genetic models, particularly in the full dataset with over 400,000 individuals and lower levels of relatedness (Tables S1-S6). Although Regenie performs better than both BLUP.QRS and QRS, it can also have elevated type I error rates (Tables 1, S1, S2, and S6). For instance, Regenie reaches a type I error rate of 9.74 *×* 10^−6^ at significance level of 10^−6^ under the heterogeneous model (Table 1), and 1.22 *×* 10^−7^ at significance level of 10^−8^ under the *t*_(2)_ model (Table S1). In contrast, Regenie.QRS consistently maintains well-calibrated type I error rates across all genetic models and significance thresholds. For example, empirical type I error rates for Regenie.QRS remain close to or below the nominal level of 10^−6^ across quantiles and models (Tables 1 and 2). Minor deviations are observed only under the particularly challenging *t*_(2)_ error model with over 400,000 samples at the most stringent thresholds (10^−8^ and 10^−7^), where all methods exhibit some inflation (Table S2). Notably, under the extreme relatedness scenario with 46,887 first-degree relatives, Regenie.QRS is the only method that controls type I error within the 95% confidence interval across significance levels from 10^−4^ to 10^−8^ (Tables S5-S8). Overall, these results highlight Regenie.QRS as the most reliable and robust method for maintaining type I error control across diverse simulation settings.

**Table 1:** Empirical type I error rates at significance level 10^−6^ with 409,627 white British individuals across all models. Note that < 5.84 *×* 10^−8^ indicates that no single SNP among the 171,159 null SNPs was found to be significant across the 100 replications.

|  |  | Homogenous | t2 | Dominance | Heterogenous |
| --- | --- | --- | --- | --- | --- |
|  | Quantiles | Empirical type I error |  |  |  |
| Regenie | - | $1.46 \times 10^{-6}$ | $2.92 \times 10^{-6}$ | $1.22 \times 10^{-6}$ | $9.74 \times 10^{-6}$ |
| QRS | composite | $5.48 \times 10^{-6}$ | $1.49 \times 10^{-5}$ | $4.26 \times 10^{-6}$ | $3.05 \times 10^{-6}$ |
| | 0.2 | $3.41 \times 10^{-6}$ | $1.22 \times 10^{-5}$ | $1.22 \times 10^{-6}$ | $1.83 \times 10^{-6}$ |
| | 0.5 | $5.48 \times 10^{-6}$ | $1.37 \times 10^{-5}$ | $3.05 \times 10^{-6}$ | $4.26 \times 10^{-6}$ |
| | 0.8 | $3.84 \times 10^{-6}$ | $7.73 \times 10^{-6}$ | $3.05 \times 10^{-6}$ | $2.44 \times 10^{-6}$ |
| Regenie.QRS | composite | $1.34 \times 10^{-6}$ | $3.53 \times 10^{-6}$ | $1.22 \times 10^{-6}$ | $2.44 \times 10^{-6}$ |
| | 0.2 | $9.74 \times 10^{-7}$ | $1.95 \times 10^{-6}$ | $1.83 \times 10^{-6}$ | $< 5.84 \times 10^{-8}$ |
| | 0.5 | $1.58 \times 10^{-6}$ | $3.23 \times 10^{-6}$ | $1.22 \times 10^{-6}$ | $1.83 \times 10^{-6}$ |
| | 0.8 | $9.74 \times 10^{-7}$ | $1.71 \times 10^{-6}$ | $1.22 \times 10^{-6}$ | $1.83 \times 10^{-6}$ |
| BLUP.QRS | composite | $3.05 \times 10^{-6}$ | $5.24 \times 10^{-6}$ | $4.87 \times 10^{-6}$ | $2.44 \times 10^{-6}$ |
| | 0.2 | $2.80 \times 10^{-6}$ | $3.53 \times 10^{-6}$ | $3.65 \times 10^{-6}$ | $1.83 \times 10^{-6}$ |
| | 0.5 | $3.41 \times 10^{-6}$ | $4.38 \times 10^{-6}$ | $3.65 \times 10^{-6}$ | $1.22 \times 10^{-6}$ |
| | 0.8 | $2.01 \times 10^{-6}$ | $3.84 \times 10^{-6}$ | $2.44 \times 10^{-6}$ | $4.87 \times 10^{-6}$ |

**Table 2:** Empirical type I error rates at significance level 10^−6^ with 46,887 first-degree white British relatives across all models.

|  |  | Homogenous | t2 | Dominance | Heterogenous |
| --- | --- | --- | --- | --- | --- |
|  | Quantiles | Empirical type I error |  |  |  |
| Regenie | - | $6.43 \times 10^{-6}$ | $1.44 \times 10^{-5}$ | $1.40 \times 10^{-6}$ | $1.11 \times 10^{-6}$ |
| QRS | composite | $3.27 \times 10^{-6}$ | $4.44 \times 10^{-6}$ | $4.50 \times 10^{-6}$ | $2.75 \times 10^{-6}$ |
| | 0.2 | $3.15 \times 10^{-6}$ | $4.50 \times 10^{-6}$ | $3.10 \times 10^{-6}$ | $1.93 \times 10^{-6}$ |
| | 0.5 | $3.51 \times 10^{-6}$ | $5.02 \times 10^{-6}$ | $4.21 \times 10^{-6}$ | $2.51 \times 10^{-6}$ |
| | 0.8 | $2.86 \times 10^{-6}$ | $3.97 \times 10^{-6}$ | $3.04 \times 10^{-6}$ | $2.40 \times 10^{-6}$ |
| Regenie.QRS | composite | $5.26 \times 10^{-7}$ | $1.23 \times 10^{-6}$ | $1.23 \times 10^{-6}$ | $7.01 \times 10^{-7}$ |
| | 0.2 | $1.05 \times 10^{-6}$ | $9.93 \times 10^{-7}$ | $9.35 \times 10^{-7}$ | $1.23 \times 10^{-6}$ |
| | 0.5 | $8.76 \times 10^{-7}$ | $1.23 \times 10^{-6}$ | $8.18 \times 10^{-7}$ | $5.84 \times 10^{-7}$ |
| | 0.8 | $3.51 \times 10^{-7}$ | $1.46 \times 10^{-6}$ | $8.76 \times 10^{-7}$ | $1.93 \times 10^{-6}$ |
| BLUP.QRS | composite | $2.10 \times 10^{-6}$ | $2.51 \times 10^{-6}$ | $2.75 \times 10^{-6}$ | $2.16 \times 10^{-6}$ |
| | 0.2 | $2.40 \times 10^{-6}$ | $2.34 \times 10^{-6}$ | $1.99 \times 10^{-6}$ | $2.34 \times 10^{-6}$ |
| | 0.5 | $2.63 \times 10^{-6}$ | $2.16 \times 10^{-6}$ | $2.51 \times 10^{-6}$ | $1.64 \times 10^{-6}$ |
| | 0.8 | $2.04 \times 10^{-6}$ | $2.75 \times 10^{-6}$ | $2.04 \times 10^{-6}$ | $1.40 \times 10^{-6}$ |

Power is calculated as the proportion of significant SNPs among the causal SNPs, averaged across 100 replications. For the homogeneous setting with normal and *t*_(2)_ error models, power is evaluated among the 10,000 causal SNPs located on the odd chromosomes. Under the heterogeneous and dominance models, power is calculated based on 100 SNPs that exhibit heterogeneous or dominance effects, respectively, averaged across 100 replications.

Under the homogeneous model, Regenie.QRS exhibits similar (only slightly lower) power than Regenie (Fig. 1, Tables S9-S10). For example, with over 400,000 samples and at a significance level of 0.05, the power is 0.78 for Regenie.QRS and 0.79 for Regenie, as the assumptions of the linear model are perfectly satisfied. Regenie.QRS achieves higher power than QRS (i.e., 0.71) and BLUP.QRS (i.e., 0.73) under the homogeneous setting, with the difference being larger as the sample size increases from 46,000 to 400,000 (Fig. 1 and Tables S9-S10). Under the *t*_(2)_ error model and a significance level of 0.05, Regenie.QRS achieves similar power (0.81) as Regenie (0.80), and demonstrates a clear advantage over QRS (0.73) and BLUP.QRS (0.75). Similar results hold at smaller alpha level of 10^−6^ (Tables S11-S12).

**Figure 1:**
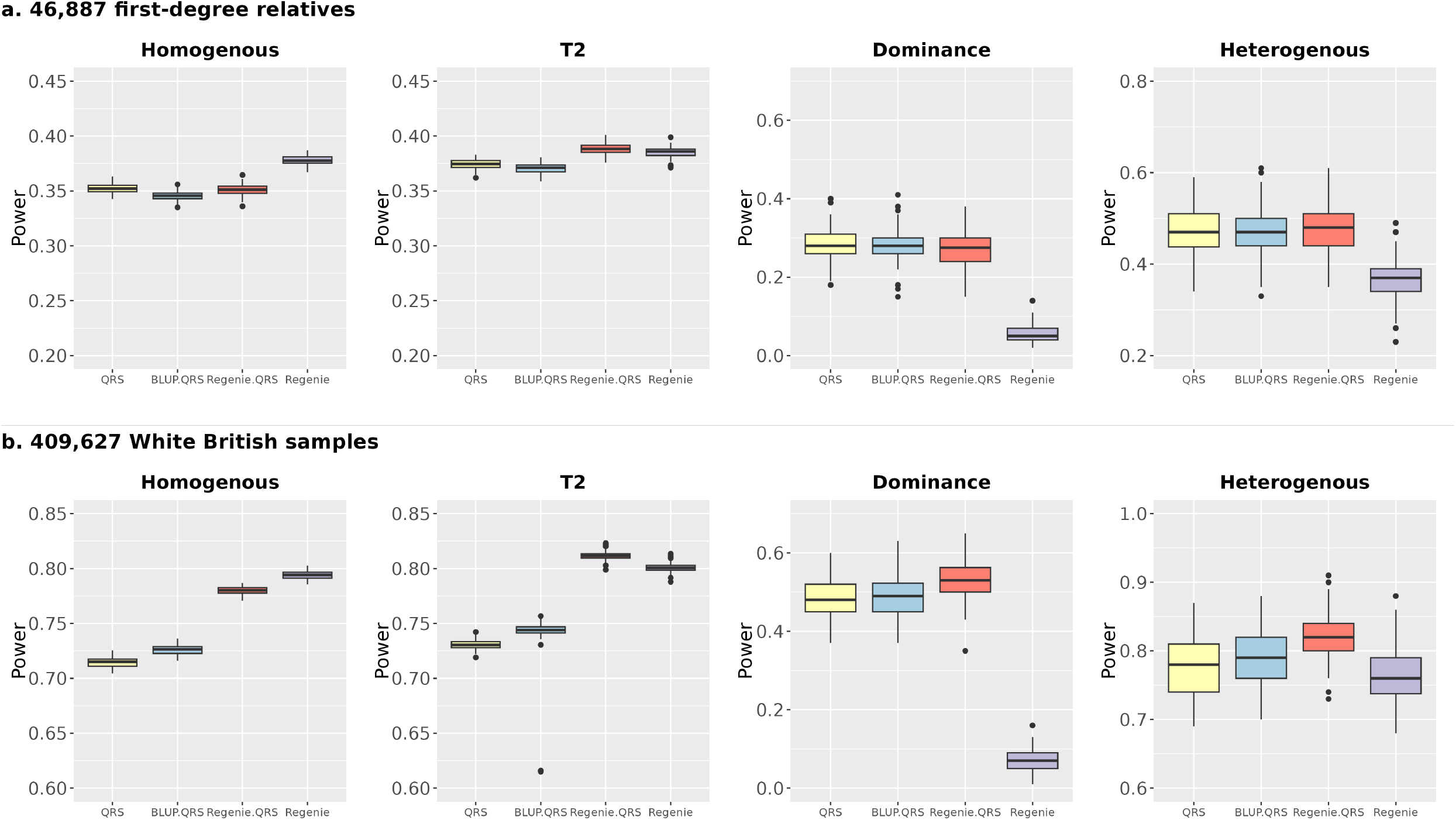
Power comparison of QRS, Regenie.QRS, BLUP.QRS, and Regenie across all model settings. Power is calculated as the proportion of causal SNPs detected at the 0.05 significance level, averaged over 100 replications; **a**. 46,887 first-degree related individuals; **b**. 409,627 White British samples.

Under the dominance model, Regenie.QRS substantially outperforms Regenie (Fig. 1, Tables S13-S14). With a sample of 46,887 first-degree relatives, Regenie.QRS achieves 21% higher power than Regenie at 0.05 significance level (Fig. 1). With a larger sample size of over 400,000, Regenie.QRS shows a 46% increase in power compared to Regenie and a 5% power improvement over both QRS and BLUP.QRS (Fig. 1). Even at a more stringent alpha level such as 10^−6^, Regenie.QRS achieves 28% power, whereas Regenie shows nearly zero power (Table S14).

Under the heterogeneous model, Regenie.QRS also demonstrates superior performance (Fig. 1, Tables S15-S16). With 46,887 first-degree relatives, Regenie.QRS achieves 10% higher power than Regenie across various significance levels (Fig. 1 and Table S15). As the sample size increases to 400,000, the power difference between Regenie.QRS and Regenie narrows at the 0.05 significance level, with Regenie.QRS maintaining a modest 6% advantage. However, at more stringent thresholds, Regenie.QRS exhibits substantially greater power, i.e. 17% and 20% higher than Regenie at the 10^−6^ and 10^−8^ alpha levels, respectively (Table S16).

For homogenous setting, we further restricted the analysis to 10,000 individuals with the highest kinship coefficients estimated by KING. FPR and power for all methods are reported in Supplemental Tables S17 and S18, respectively. Conclusions are similar to those with 46,887 samples.

### Applications to continuous traits in UKBB and ProgeNIA/SardiNIA

We first conducted Regenie.QRS analyses and compared the results with those from Regenie using UKBB imputed data from all white British participants. To perform quality control, we excluded individuals with more than 10% missing genotypes, included only SNPs with a MAF greater than or equal to 0.05, considered only SNPs with a 90% genotyping rate, and filtered out SNPs with a Hardy-Weinberg disequilibrium p-value less than 10^−4^. Missing genotypes were imputed by assigning the most common non-missing allele for each SNP using PLINK2. After quality control, we retained 382,402 individuals with 6,364,586 SNPs. For validation, we leverage an independent dataset from the ProgeNIA/SardiNIA project, a longitudinal study comprising a cohort of Sardinia subjects (*n* = 7, 926). Using the same quality control procedure, we obtained a total of *∼* 6, 150 individuals with 5,989,547 SNPs.

We perform GWAS for 28 traits that are shared between UKBB and ProgeNIA/SardiNIA project data, adjusting for covariates including age, sex, batch and top 10 PCs of genetic variation. Traits are normalized using a rank-based inverse normal transformation for Regenie and Regenie.QRS, as polygenic prediction by Regenie tends to be more accurate when applied to normalized phenotypes. We identify significant SNPs using a genome-wide significance threshold of 5 *×* 10^−8^ for each trait. To identify independent genetic loci, we performed LD clumping using PLINK2 with an LD *r*^2^ threshold of 0.1 and a clumping window of 1000 kb. Each locus was represented by its lead variant, defined as the variant with the smallest p-value within that region. We then mapped these lead variants to their nearest genes. Lead SNPs were grouped together if they were located within a 1 Mb window or mapped to the same gene. We compared the sets of loci identified by Regenie and Regenie.QRS, quantifying the number of shared and unique loci detected by each method (Fig. 2).

Overall, Regenie identifies more loci suggesting possibly higher power but also possible higher rate of false positives. Most of the Regenie.QRS discoveries are shared with Regenie, but Regenie.QRS also identifies new loci that Regenie misses. Furthermore, compared with the analysis using only 320K unrelated individuals in Wang et al. ^3^, Regenie.QRS applied to the larger set of white British participants identifies substantially more discoveries after incorporating all related samples across traits, demonstrating the benefits of incorporating related individuals (Fig. 3). See also Table S19 for simulation results.

**Figure 2:**
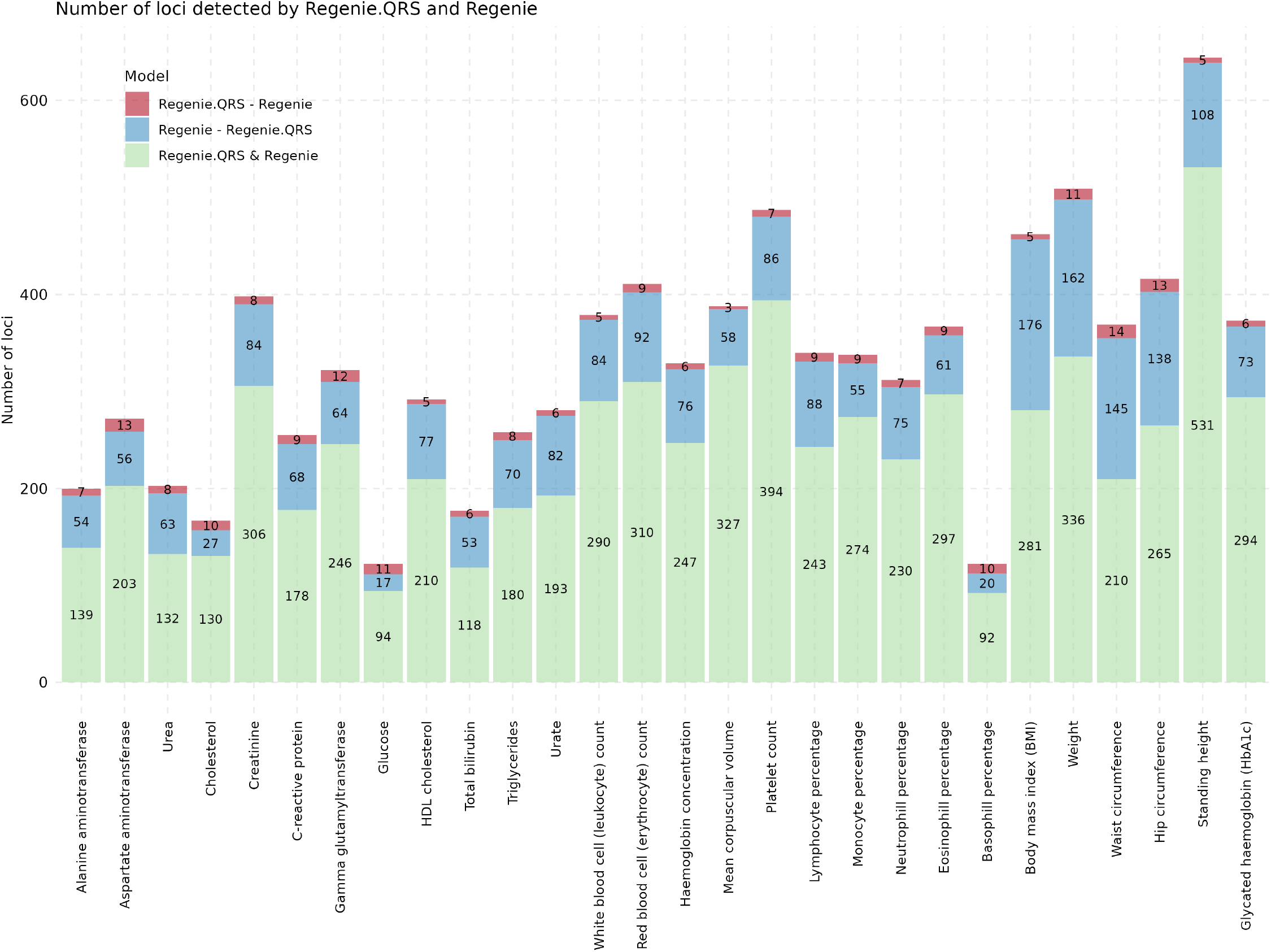
Number of genome-wide significant loci in the UK Biobank study. Stacked barplots show the corresponding number of significant loci for Regenie and Regenie.QRS. The red bars (Regenie.QRS - Regenie) indicate loci uniquely detected by Regenie.QRS but not Regenie, the blue bars (Regenie - Regenie.QRS) represent loci identified by Regenie but not Regenie.QRS, and the green bars (Regenie & Regenie.QRS) show loci commonly detected by both methods.

**Figure 3:**
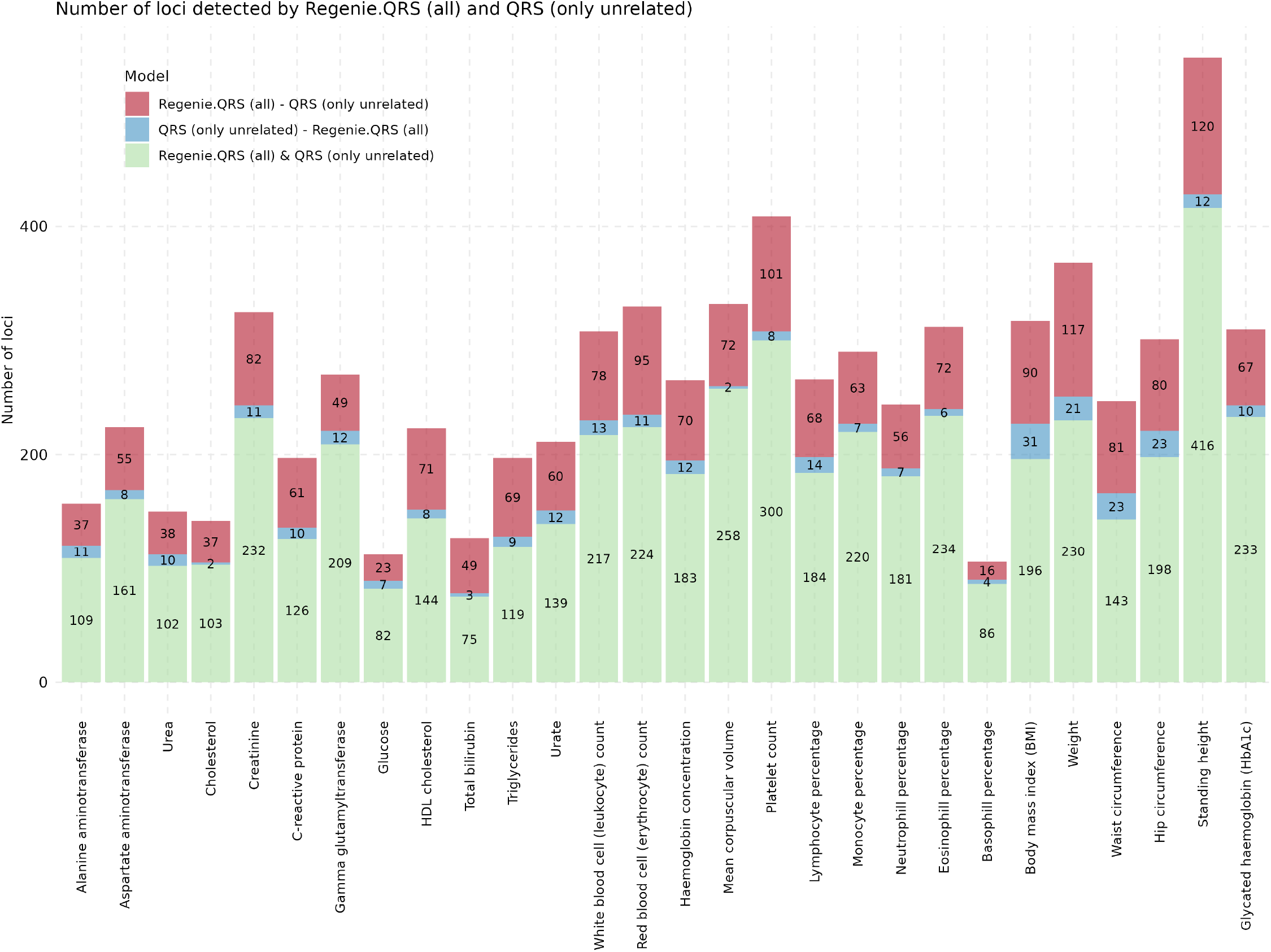
Number of genome-wide significant loci detected by QRS and Regenie.QRS. Stacked barplot shows the number of significant loci identified across traits using QRS on about 320,000 unrelated individuals in^3^ and Regenie.QRS on 382,402 individuals including related samples. Green bars represent loci detected by both methods, blue bars indicate loci uniquely detected by QRS but not Regenie.QRS, and red bars denote loci uniquely detected by Regenie.QRS but not QRS.

### Validations using the ProgeNIA/SardiNIA project data

We performed similar analyses in the ProgeNIA/SardiNIA data (Fig. S2). The results look consistent with the UKBB analysis, i.e. Regenie tends to identify more loci relative to Regenie.QRS, but Regenie.QRS also identifies loci that are missed by Regenie.

To assess the replicability of loci from the UKBB analysis in the Sardinia data, SNPs are first sorted by their UKBB p-values. The SNP with the smallest p-value in UKBB that meets the genome-wide significance threshold is selected as the lead SNP for the first clump. SNPs located within 1 Mb of the lead SNP and having an *R*^2^ *>* 0.8 are considered correlated with the lead SNP. When calculating the replication rate, we check whether any SNPs within the same clump have p-values in the Sardinia data below the 0.05 significance level. Overall, the replication rates between the two methods are generally consistent, but low given the small sample size for the replication dataset (Fig. 4).

**Figure 4:**
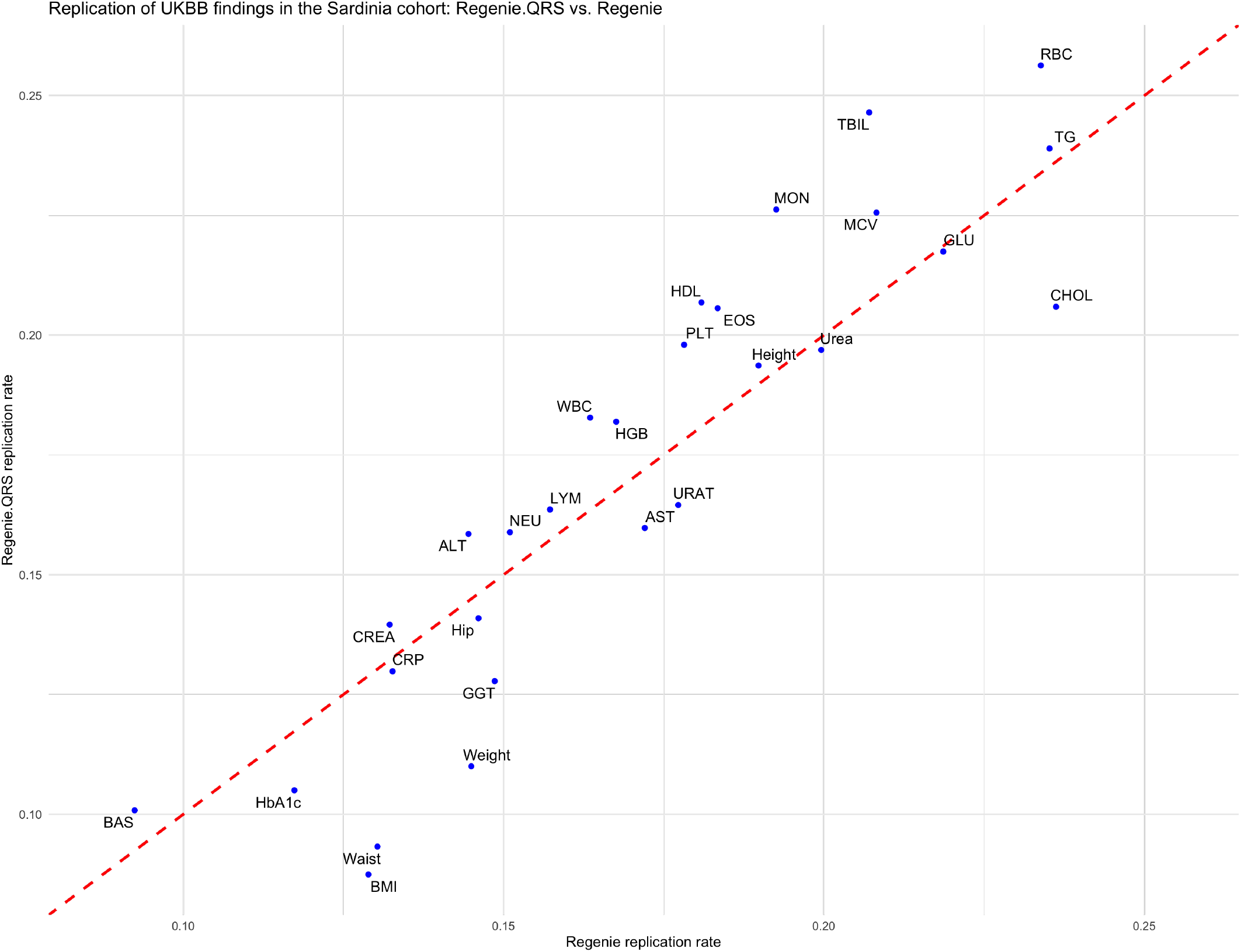
Regenie.QRS vs. Regenie replication rates. Each blue dot represents a phenotype with its replication rate from Regenie (x-axis) and the Regenie.QRS (y-axis). The red dashed line indicates the 45 degree line. Abbreviations: ALT – Alanine aminotransferase; AST – Aspartate aminotransferase; CHOL – Cholesterol; CREA – Creatinine; CRP – C-reactive protein; GGT – Gamma glutamyltransferase; GLU – Glucose; HDL – HDL cholesterol; TBIL – Total bilirubin; TG – Triglycerides; URAT – Urate; WBC – White blood cell count; RBC – Red blood cell count; HGB – Haemoglobin concentration; MCV – Mean corpuscular volume; PLT – Platelet count; LYM – Lymphocyte percentage; MON – Monocyte percentage; NEU – Neutrophil percentage; EOS – Eosinophil percentage; BAS – Basophil percentage; BMI – Body mass index; Waist – Waist circumference; Hip – Hip circumference; Height – Standing height; HbA1c – Glycated haemoglobin.

### Examples of heterogeneous effects

One of the main advantages of quantile regression is that it allows a more detailed characterization of genetic effects across different parts of the phenotype distribution. We illustrate here several examples where genetic variants have heterogeneous or varying effects across phenotypic quantiles (additional examples are presented in the Supplemental Material). These loci are identified by both Regenie.QRS and Regenie, and replicated in the Sardinia analysis. The lead SNPs at these loci exhibit heterogeneous effect sizes across quantile levels.

#### G6PC2 and glucose levels

An interesting example is for glucose level. We identified a highly significant association at the *G6PC2* locus, with lead QR SNP rs13431652 located in an enhancer for *G6PC2* (Fig. 5). This variant shows highly significant associations in both the UKBB and Sardinia cohorts. Previous studies have identified rs13431652 as strongly associated with elevated fasting glucose^33^, and nominated it as a likely causal regulatory variant by statistical fine-mapping and functional experiments^34;35^. Functionally, the major allele T enhances binding of the NF-Y transcription factor, increasing *G6PC2* promoter activity in pancreatic islets and contributing to elevated fasting glucose via modulation of *β*-cell glycolytic flux ^33^. Our quantile regression analysis also detects heterogeneous effects for the major allele T with strongest effect sizes at lower quantiles of glucose distribution (i.e., *β ≈* 0.10 at quantile 0.1), and significantly lower effects for the higher quantiles (i.e., *β ≈* 0.02 – 0.03 at quantile 0.9). Note that, in contrast, Regenie only provides a single effect size estimate (0.088), which can be interpreted as an average effect across the different quantiles. This pattern suggests that the physiological impact of rs13431652 is more pronounced in individuals with lower glucose levels, acting as a guardian against low glucose without driving dangereous hyperglycemia. This likely explains the lack of association of this variant in GWAS for type 2 diabetes^33;34^. This trend is replicated in the Sardinia cohort, where effect sizes remain positive ( *≈* 0.05 – 0.075) but confidence intervals are wider due to smaller sample size.

**Figure 5:**
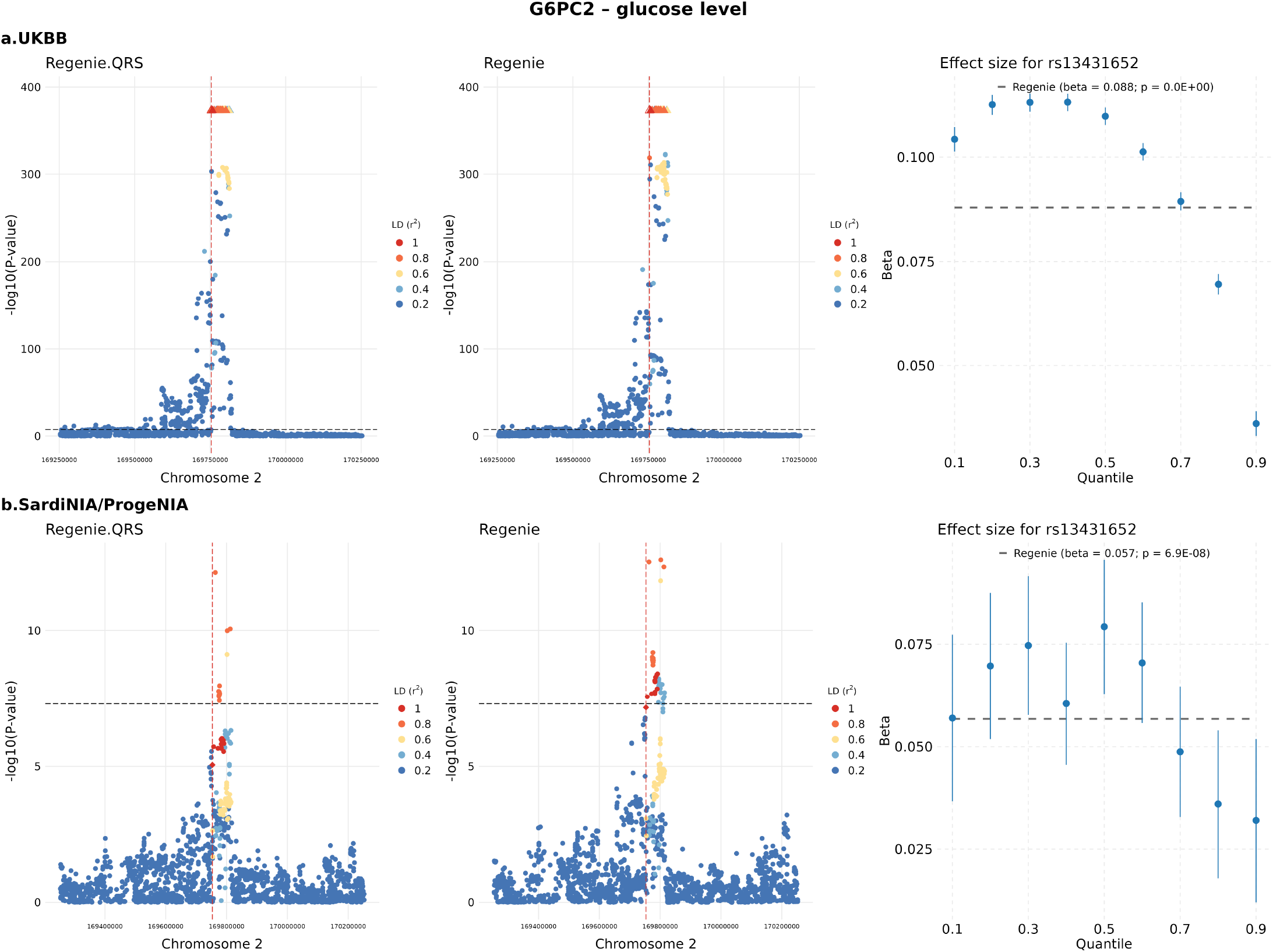
Heterogeneous effect of rs13431652 located at the *G6PC2* locus and glucose levels. Locus zoom plots display −log_10_(p-value) for each variant, with the lead SNP at each locus shown as a diamond and highlighted by a vertical dashed line. Colors indicate the linkage disequilibrium (LD, *r*^2^) between each variant and the lead SNP. **a**. UKBB; **b**. Sardinia. The horizontal dashed line marks the genome-wide significance threshold at 5 *×* 10^−8^. For the lead Regenie.QRS SNP, the quantile-specific effect sizes *±* standard errors are shown in the right panel at different quantile levels.

#### SORT1 and total cholesterol

For total cholesterol, one of the strongest association signals emerges at the *SORT1* locus on chromosome 1, with lead SNP rs12740374 (Fig. 6). This SNP has been previously identified as a causal variant contributing to change in plasma LDL-C ^36^. This association is highly significant in UKBB ( −log_10_ *p >* 200) in both Regenie and Regenie.QRS and is replicated in the Sardinia cohort. rs12740374 is a noncoding variant which affects cholesterol metabolism by creating a C/EBP transcription factor binding, increasing *SORT1* expression in the liver and leading to decreases in plasma LDL-C levels ^37^. Moreover, this SNP has been shown to influence pharmacogenetic response to statin therapy. Individuals carrying the minor allele T exhibit greater reductions in LDL-C compared to non-carriers when treated, suggesting that SNP rs12740374 modulates the efficacy of lipid-lowering interventions^37^. Quantile regression reveals a clear increasing protective effect of the minor allele T across cholesterol quantiles in UKBB, with stronger effects observed at the upper tails of the distribution, which highlights quantile-dependent heterogeneity that is not possible to characterize using LR models like Regenie. In the Sardinia cohort, effect sizes for the T allele fall within a similar range (-0.03 to -0.09) and follow the same direction but with wider confidence intervals due to small sample size. The fact that the most protective effect is at the upper quantiles, the highest risk group from a clinical standpoint, and that statins are more effective in reducing LDL cholesterol levels in individuals with the T allele, can have important consequences for drug dosage and drug development.

**Figure 6:**
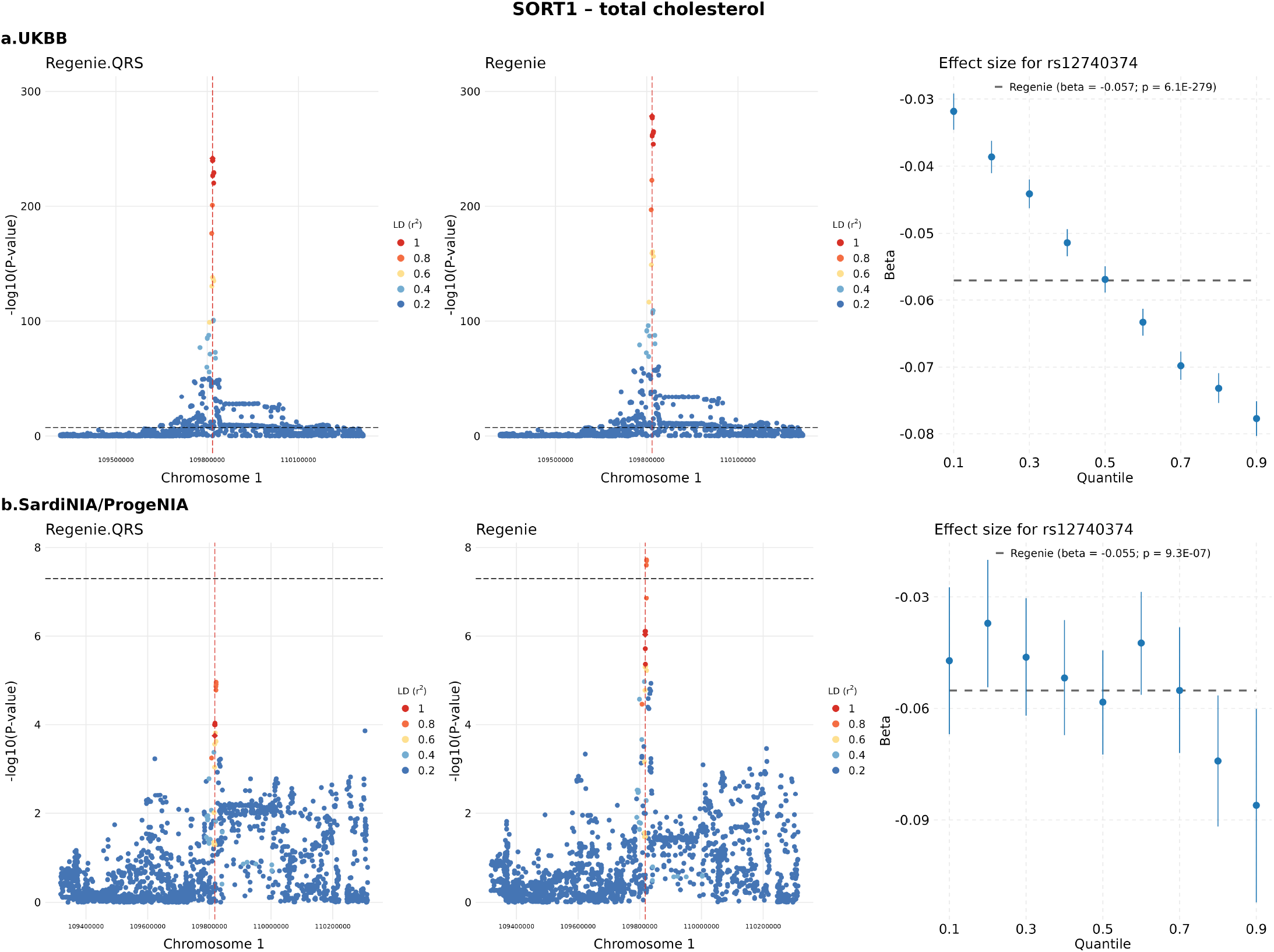
Heterogeneous effect of rs12740374 located at the *SORT1* locus and total cholesterol. Locus zoom plots display −log_10_(p-value) for each variant, with the lead SNP at each locus shown as a diamond and highlighted by a vertical dashed line. Colors indicate the linkage disequilibrium (LD, *r*^2^) between each variant and the lead SNP. **a**. UKBB; **b**. Sardinia. The horizontal dashed line marks the genome-wide significance threshold at 5 *×* 10^−8^. For the lead Regenie.QRS SNP, the quantile-specific effect sizes *±* standard errors are shown in the right panel at different quantile levels.

### Computational performance

We evaluated the computational performance of Regenie.QRS and BLUP.QRS by considering both memory usage and projected CPU time (Table S20). The time complexity for Step 1 polygenic predictions for Regenie.QRS and Regenie.BLUP model are the same as in Regenie ^14^ and SAIGE^28^, respectively. Regenie achieves faster polygenic prediction compared to BLUP in SAIGE by operating on block-wise genotype matrices^14^. For similar reasons it is also more memory efficient. Regenie.QRS (and BLUP.QRS) will incur higher computational cost than Regenie (and SAIGE) due to the inherent increase in computation for quantile regression testing vs. linear regression testing in the second step.

We show a concrete example from the application to BMI, for 382,402 individuals and 6,364,586 variants (Table S20).

#### Polygenic prediction

Polygenic prediction using Regenie was completed in 8 CPU hours, while the BLUP prediction using SAIGE took approximately 7 hours on 4 CPU cores (i.e. 28 CPU hours).

#### Association testing

##### Regenie

Step 2 association analyses with Regenie were conducted independently for each chromosome in parallel, requiring approximately 1.5 CPU hours per chromosome (i.e. 33 CPU hours).

##### QRS

QRS testing required approximately 1,144 CPU hours in total.

##### Regenie.QRS and BLUP.QRS

After generating polygenic predictions by Regenie, Regenie.QRS tests required 652 CPU hours in total. Similarly, BLUP.QRS testing required approximately 571 CPU hours.

Note that QRS, Regenie.QRS, and BLUP.QRS can all be significantly accelerated by parallelizing tests across small genomic regions, and therefore the actual elapsed wall clock time is much shorter. Specifically, we divided the genome-wide variant set into 1,073 segments, each containing approximately 6,000 variants. After generating polygenic predictions using Regenie, the elapsed wall-clock time was reduced to 36 minutes with 1,073 CPU cores. Similarly, BLUP.QRS took 32 minutes per segment, while QRS required about 1.1 hours per segment.

## Discussion

We presented a computationally efficient whole-genome quantile regression framework for quantitative traits in biobank scale GWAS studies. Regenie.QRS provides a counterpart of the linear regression method Regenie in the context of quantile regression, and extends existing marginal quantile regression models to whole-genome regression. Our applications underscore the computational efficiency and robustness of Regenie.QRS in uncovering complex genotype-phenotype interactions, particularly for traits characterized by non-normal distributions and heterogeneous effects. A main appeal of Regenie.QRS is its ability to provide a more complete picture of the genetic effects across different parts of the distribution which can pave the way for more in-depth mechanistic investigations.

A possible limitation of quantile regression is that it requires the availability of individual level data and cannot leverage existing summary statistics. However, with the increasing trend in molecular and biomarker data generation across many studies, quantile regression methods as proposed here will only increase in relevance. Furthermore, recent developments in secure federated genome-wide association studies allow for the application of such methods to private data held by multiple parties while maintaining data confidentiality ^38^.

Another possible limitation is that quantile regression tends to identify fewer loci than linear regression in applications which likely reflects higher power for the linear regression model, although some false positive inflation can also account for some of the discoveries. We argue that in the current era where GWAS findings reach saturation for many traits, and when the goal moves from discovery to mechanistic understanding of existing discoveries, more interpretable models such as proposed here are highly valuable.

An implicit assumption in our framework is that the GRM does not vary across quantiles. In practice, the genetic architecture of complex traits may exhibit differential effects at different quantiles, implying that the structure of the GRM itself could be quantile-dependent. Future extensions of this work could explore adaptive GRM estimation tailored to specific quantiles, further improving power in detecting heterogenous genetic effects.

Beyond genetic discovery, QR can have interesting applications to genomic trait prediction. In the typical polygenic risk score (PRS) prediction task the aim is to predict the mean of a phenotype (*Y* ) given the genotype profile of an individual (*G*). The resulting PRS is a linear combination of individual genotypes weighted by estimated genetic effects from a linear regression model in GWAS. This is essentially a point prediction, meant to reflect the typical or average phenotypic outcome given a particular genotype profile, i.e. *E*(*Y* | *G*). The appeal of QR is that it extends mean-based LR to the analysis of the entire conditional distribution of a phenotype of interest *Q*_*Y*_ (*τ* | *G*) for any quantile level *τ ∈* (0, 1). Thus QR allows us to move away from point predictions to prediction intervals that naturally account for the uncertainty in the point estimation. This goal will require the scalable implementation of penalized regression methods^39^, such as elastic net combined with a quantile loss function to ensure genome-wide scalability and accuracy^40^. These advancements have the potential to greatly expand the applicability of quantile regression for high-dimensional genomic data, providing powerful tools to better understand complex genetic architectures. We leave these extensions and developments for future research.

## Acknowledgements

F.W. and I.I.L. are supported by NIH grants MH095797 and AG072272. This research has been conducted using the UK Biobank Resource under Application Number 41849.

## Data Availability

The individual-level genotype and phenotype data are available to approved researchers through the UK Biobank web portal at https://www.ukbiobank.ac.uk/. Regenie.QRS GWAS summary statistics for the biomarkers reported here are publicly available at https://doi.org/10.5281/zenodo.11095249.

## Code Availability

Regenie.QRS and other QR scripts: https://github.com/Iuliana-Ionita-Laza/QRGWAS. Regenie: https://rgcgithub.github.io/Regenie/.

## Supplemental Material

### Asymptotic properties

In this section, we demonstrate the consistency of the Regenie and BLUP estimators of polygenic effects, and the asymptotic linear representation of QR estimation, accounting for polygenic predictions.

For Regenie, let *ĝ*_*i*,*k*_ represent the first-stage polygenic prediction by ridge regression within block *k* for the *i*-th individual. The second-level ridge regression takes the linear combination of the first-step ridge predictions, denoted by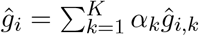.

#### Theorem 1

*Under regularity conditions that ridge estimator is consistent in the L*_2_ *norm* ^*41*^, *assuming* 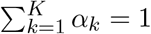 *and* 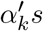 *are non-negative, then polygenic predictions from Regenie ĝ*_*i*_ *satisfy* 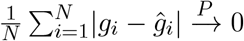.

**Proof**. Assuming that the ridge estimator is consistent in the *L*_2_ norm ^41^, consistency in the *L*_1_ norm can be achieved by applying Hölder’s inequality. Specifically, we have:

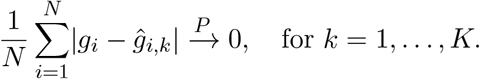

Then in the second-level ridge regression,

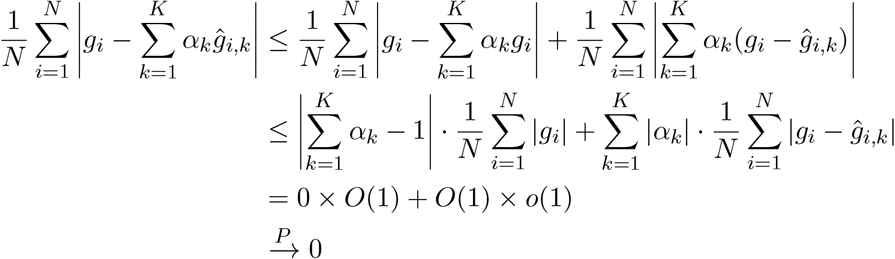

Although Theorem 1 holds only under the condition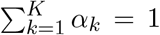, simulations in ^23^ have shown that this constraint is largely unnecessary. The decreases in prediction error are almost identical whether or not the constraint 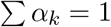 is used ^23^.

#### Theorem 2

*Assume the regularity conditions stated in Theorem 3.2 of*^*42*^ *hold, and ∥***QΛ**^1*/*2^∥_1,*∞*_ *≤ L, where L is a constant. Then the BLUP in* (6) *satisfies*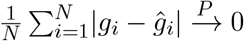.

**Proof**. Jiang et al. ^42^ has derived the asymptotic results for BLUPs for random slopes whose components are i.i.d. We define *g*^*\*\**^ to be Λ^−1*/*2^*Q*^⊤^***g***, where *Q* and Λ denote the eigenvectors and eigenvalues of *g*, respectively. As *Q* and Λ depend on the sample size *N* , we assume that ∥*Q*Λ^1*/*2^ ∥_1, *∞*_ is bounded by *L* such that the maximum row sum of the absolute values of the matrix entries do not grow with *N* . Note that *g*^*\*\**^ has i.i.d. components with variance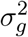. In the proof of theorem 3.2^42^, it has been shown that

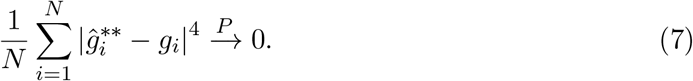

Therefore,

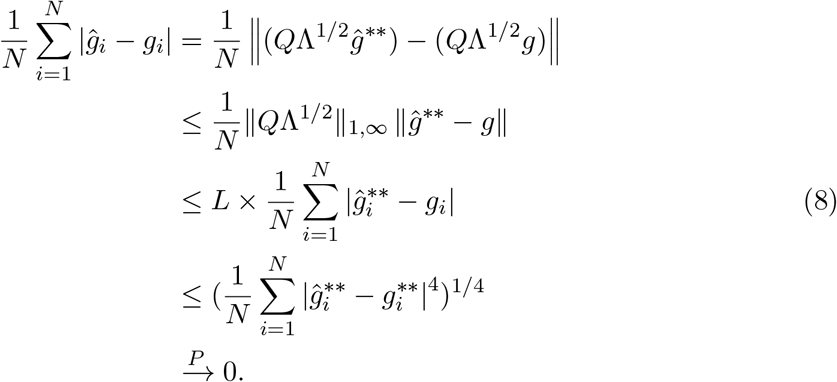

The inequality in (8) is obtained by the power mean inequality.

While computing the BLUP in (6) (Methods), the terms **Σ**^−1^***Y*** and **Σ**^−1^***Z*** pose significant computational bottlenecks when **Σ** is large and dense. To address this, we employ the Preconditioned Conjugate Gradient (PCG) method to efficiently approximate expressions of the form **Σ**^−1^***u*** for vectors ***u*** *∈* ℝ^*N*^ . Based on the error bound for the *k*th iteration provided in ^43^, it can be shown that 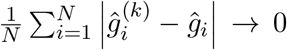 as *k → ∞*, where the 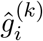 represents the approximation of BLUP at *k*-th iteration. In combination with Theorem 2 and the triangle inequality, we can show that the BLUP computed via PCG is consistent (i.e. 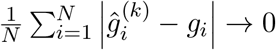 as *k → ∞*).

Theorems 1 and 2 establish the consistency of polygenic predictions obtained through Regenie and BLUP, respectively, showing that the prediction errors converge in probability to zero under their respective regularity conditions, making them reliable methods for accurate genetic prediction. Notably, the condition 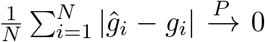 is weaker than the typical convergence requirements in the *L*_2_ or *L*_*∞*_ norms often assumed for consistent estimators. Nevertheless, it is sufficient to establish the consistency of the quantile regression estimator as shown in the next theorem. By similar argument, this condition can be satisfied by various classes of estimators used in polygenic prediction, such as linear or generalized linear lasso estimators ^44^, method of simulated moments (MSM) estimators^45^, and others.

We next show the consistency of the loss function based on polygenic predictions and the estimator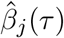. Recall that the estimation of *β*_*j*_ for SNP *j* is obtained by minimizing the quantile loss function in (3), which is 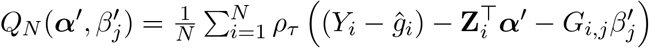, and 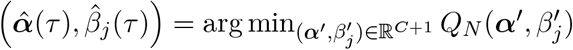. We define

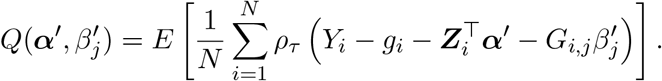

#### Theorem 3

*Under the condition*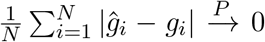 *and the regularity conditions for minimizers of convex functions*^*46*^ , *we have:*

1. 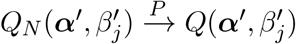.
2. 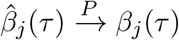.

**Proof**. When the error *ϵ* is independent and identically distributed (iid), based on the property of pinball loss function, the underlying *α*(*τ* ) and *β*_*j*_(*τ* ) in model (2) satisfy

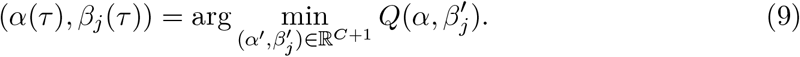

Let *ĝ* = *g* + (*ĝ − g*). Since *ρ*_*τ*_ is a Lipschitz continuous function, there exists a constant *K*_*τ*_ such that

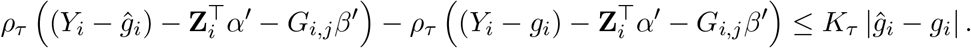

Under the condition that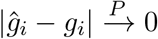,

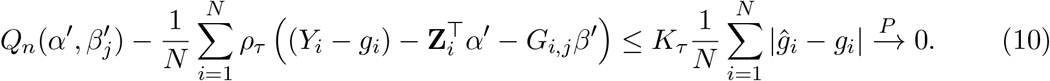

By applying the weak law of large number,

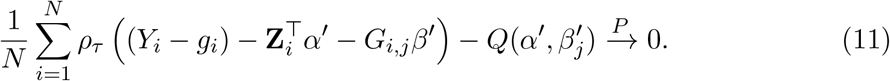

Therefore,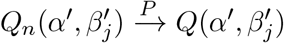 . Since 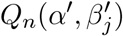 and 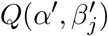 are convex functions of *α*^*′*^ and 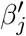, and based on the eqn. (9), we can apply the regularity conditions stated in Theorem 2.3 of ^46^ for the consistency of minimizers of convex functions. As a result, we Obtain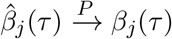.

By combining Theorem 3 with either Theorem 1 or Theorem 2, we conclude that the estimation of quantile regression which accounts for polygenic predictions from Regenie and BLUP is consistent.

#### Theorem 4

*Let ξ*_*i*_(*τ* ) = *g*_*i*_ + **Z**_**i**_^⊤^***α*(*τ* )** + *G*_*i*,*j*_*β*_*j*_(*τ* ), *and we define* 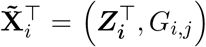 *and* ***γ*(*τ* )** = (***α*(*τ* )**, *β*_*j*_(*τ* )). *Under conditions A1–A3*^*47*^, *the asymptotic linear representation of the quantile regression estimator γ*(*τ* ) *is given by:*

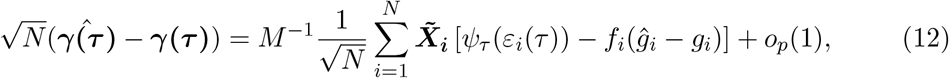

where *ψ*_τ_ (*u*) = *τ−* **1** *{u <* 0*}* denotes the quantile score function, and *f*_*i*_ is the conditional density of the error *ε*_*i*_(*τ* ) evaluated at 0 given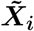.

**Proof**. We define

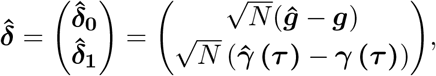

and

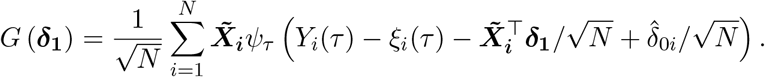

Using a similar approach and conditions as in^47^, uniformly for ∥***δ***_**1**_∥ *<* Δ_1_, one can show that

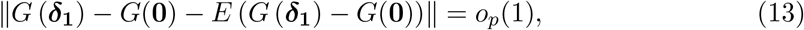

and at the minimizer, 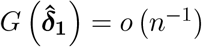. Then

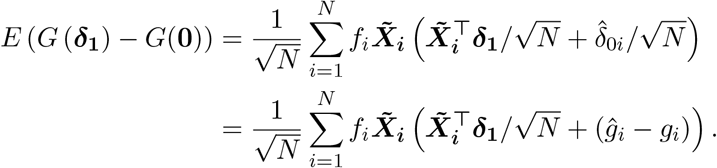

Note that 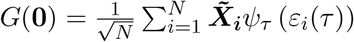 and based on eqn. (13), we could obtain

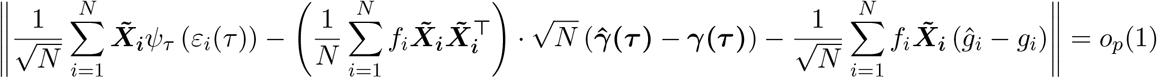

let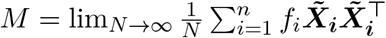, then

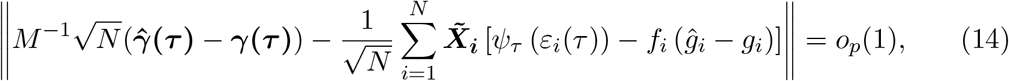

and we can obtain

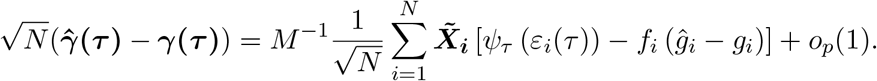

Theorem 4 establishes the asymptotic linear representation of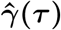, which is applicable to any type of polygenic predictions within this framework. The term *ĝ*_*i*_ can take any explicit form, and by substituting it into eqn. (12), one can derive the corresponding asymptotic form of the QR estimation. It is worth noting that, in the extreme case where the random effect is perfectly estimated (***ĝ*** = ***g***), eqn. (12) is reduced to

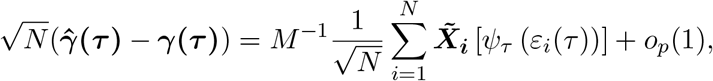

where 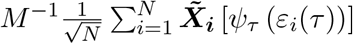 is the asymptotic linear representation. As the error 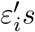 are independent and identically distributed (iid), by the central limit theorem,

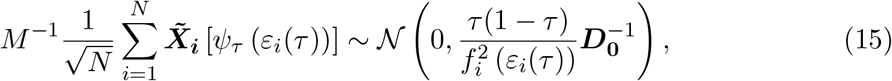

where 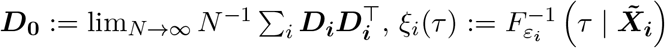, and *f* (*·*) represents the density function. The formula in (15) represents the asymptotic distribution of the classical QR estimator under i.i.d. error models^48^.

### Additional examples of loci with heterogenous effects

#### SMARCA4

A second locus for total cholesterol is tagged by the lead QR SNP rs71174352, which maps to the *SMARCA4* region on chromosome 19 (Fig. S3). *SMARCA4*, a core component of the SWI/SNF chromatin remodeling complex, plays a critical role in cholesterol regulation by modulating the accessibility of genes involved in cholesterol biosynthesis and uptake ^49^. Prior studies have linked this gene to lipid traits, including total cholesterol ^50^ and LDL cholesterol ^51^, as well as to cardiovascular outcomes such as coronary artery disease ^52^ and myocardial infarction^53^. These associations highlight SMARCA4 as a biologically plausible candidate gene for lipid regulation Regenie.QRS reveals an effect-size gradient across quantiles, uncovering heterogeneity not captured by standard association methods. Specifically, the G allele of rs71174352 demonstrates stronger effects at higher quantiles, and the pattern is replicated in both cohorts. In UKBB, effect sizes rose steadily from *∼*0.04 at quantile 0.1 to *∼*0.10 at quantile *∼*0.9 with narrow confidence intervals. The Sardinia cohort mirrored this trend despite its smaller sample size, with effects ranging from near-zero to ~0.09 at the upper tail. These consistent and quantile-dependent associations across independent populations emphasize the presence of effect heterogeneity and reinforce the functional relevance of this locus in cholesterol metabolism.

#### ATG16L1

A notable locus identified by both Regenie.QRS and Regenie for the trait total bilirubin is centered around the lead SNP rs62192912 on chromosome 2 near gene *ATG16L1* (Fig. S4). This region exhibits a dense cluster of associated variants, with many SNPs surpassing a -log10(p) value of 300. *ATG16L1* plays a key role in autophagy, a cellular process essential for maintaining liver homeostasis^54^. Emerging evidence suggests that genetic variation in *ATG16L1* is significantly associated with serum bilirubin levels^55^, potentially linking autophagy to bilirubin metabolism. Notably, rs62192912 has also been linked to total cholesterol levels in prior studies^50^, suggesting pleiotropic effects. In the Sardinia cohort analysis, this locus consistently reached genome-wide significance across both methods, with effect sizes ranging from 0.04 to 0.17. Importantly, both UKBB and Sardinia analysis demonstrated increasing effect sizes in the upper quantiles, highlighting a quantile-specific genetic architecture.

#### MPL

For triglycerides, we identified a lead SNP rs286 previously reported as significantly associated with triglyceride levels in diverse populations^56^. In our study, this locus was robustly replicated in the Sardinia cohort, with both Regenie.QRS and Regenie analyses detecting signals that exceeded the genome-wide significance threshold (Fig. S5). Notably, quantile-specific effect size analyses revealed a consistent trend across both UKBB and Sardinia datasets: the effect of rs286 increases steadily at higher quantiles, indicating a stronger genetic impact among individuals with elevated triglyceride levels.

## Supplemental Figures and Tables

**Figure S1:**
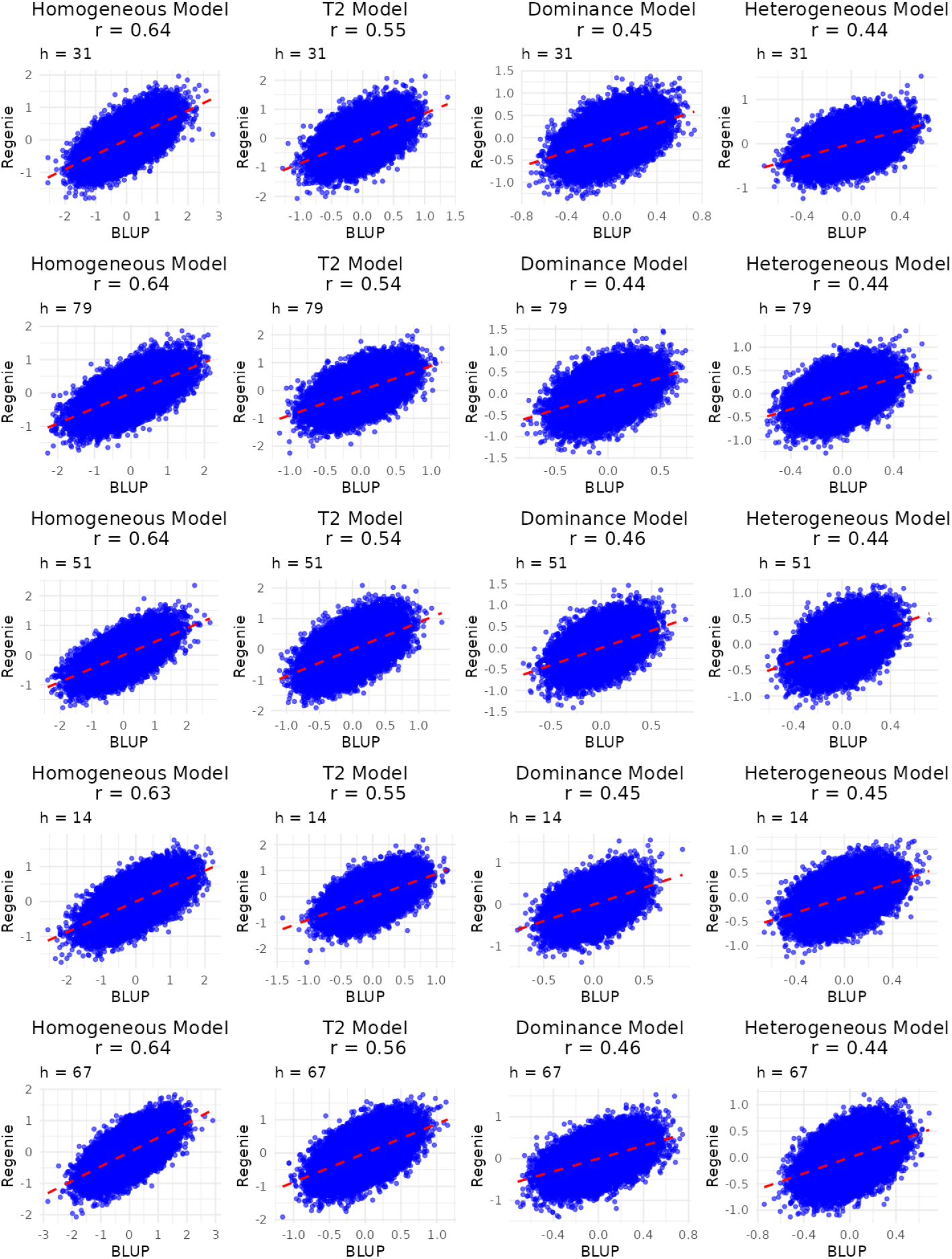
Comparison of polygenic predictions from BLUP and Regenie for chromosome 1 using LOCO scheme. Scatter plots display the relationship between BLUP (x-axis) and Regenie (y-axis) predictions across four models: Homogeneous, T2, Dominance, and Heterogeneous. Each row represents results for a randomly selected replication *h*. The Pearson correlation coefficient (*r*) is provided in each plot to quantify the agreement between the two methods. The red dashed line represents the identity line (y = x).

**Figure S2:**
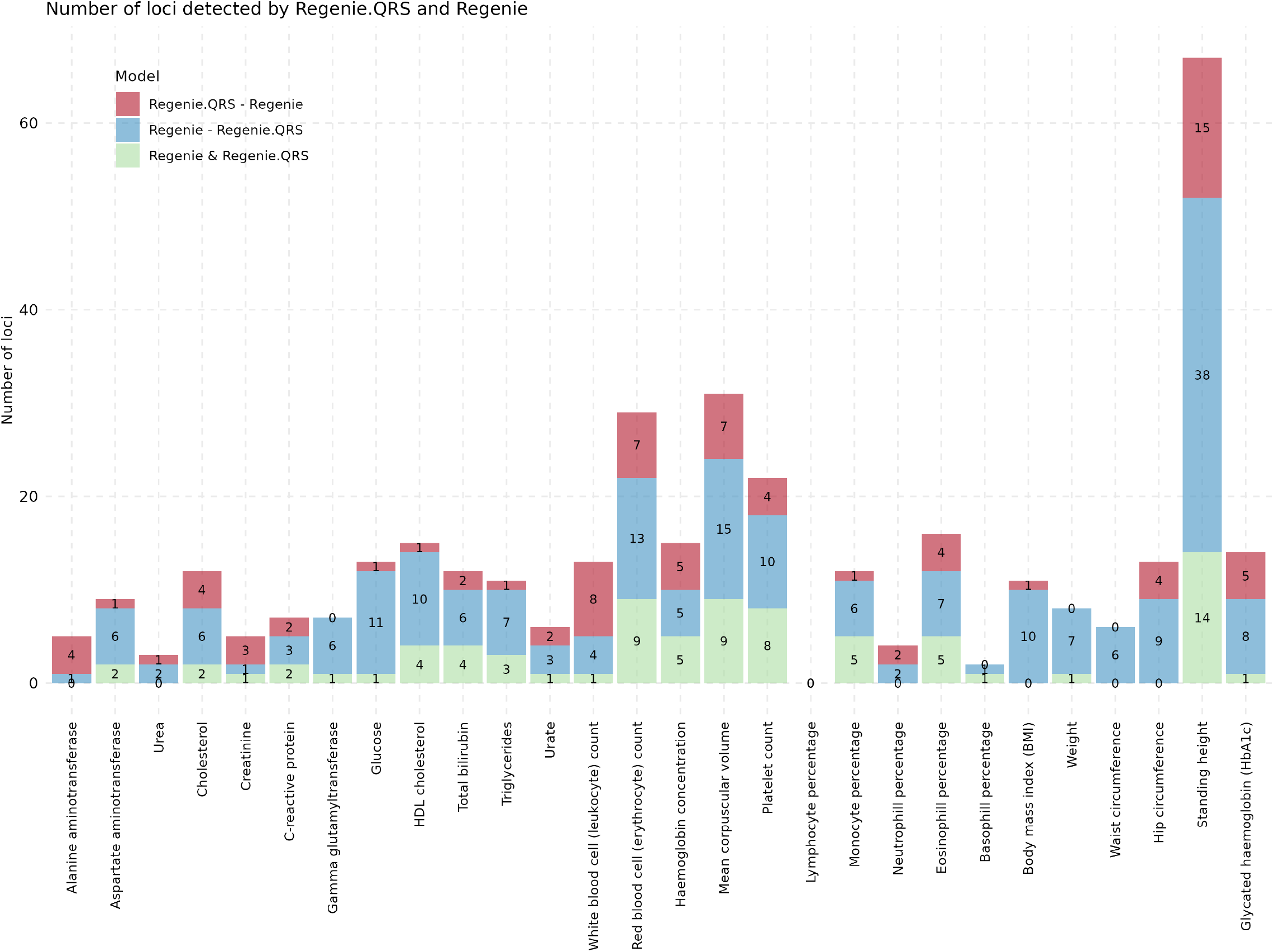
Number of significant loci at the significance level 10^−6^ in the Sardinia analysis. Stacked barplots show the corresponding number of significant loci for Regenie and Regenie.QRS. The red bars indicate loci uniquely detected by Regenie.QRS but not by Regenie, the blue bars represent loci identified by Regenie but not by Regenie.QRS, and the green bars (Regenie & Regenie.QRS) show loci commonly detected by both methods.

**Figure S3:**
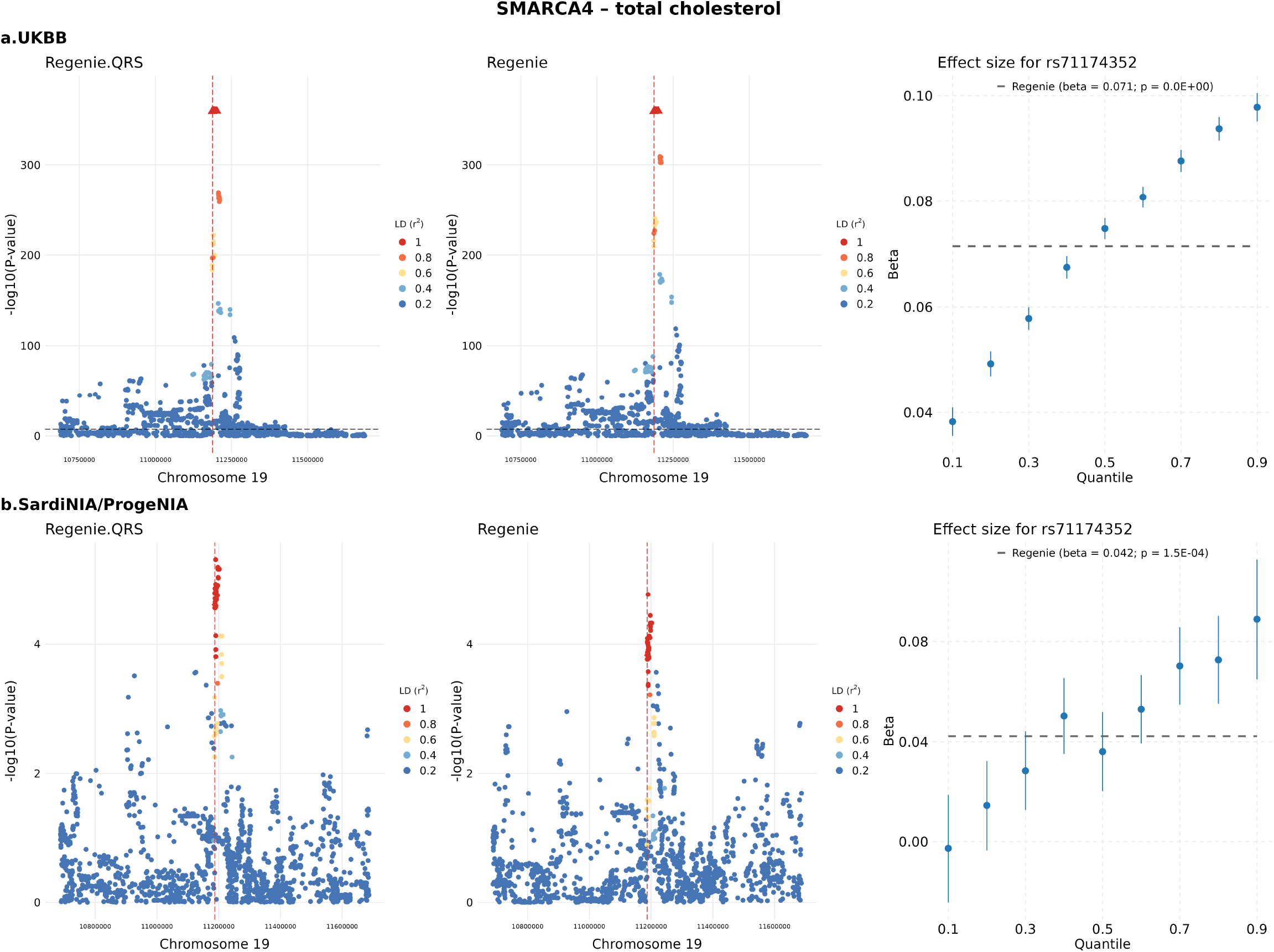
Heterogeneous effect of rs71174352 located at the *SMARCA4* locus and total cholesterol. Locus zoom plots display − log_10_(p-value) for each variant, with the lead SNP at each locus shown as a diamond and highlighted by a vertical dashed line. Colors indicate the linkage disequilibrium (LD, *r*^2^) between each variant and the lead SNP. **a**. UKBB; **b**. Sardinia. The horizontal dashed line marks the genome-wide significance threshold at 5 *×* 10^−8^. For the lead Regenie.QRS SNP, the quantile-specific effect sizes *±* standard errors are shown in the right panel at different quantile levels.

**Figure S4:**
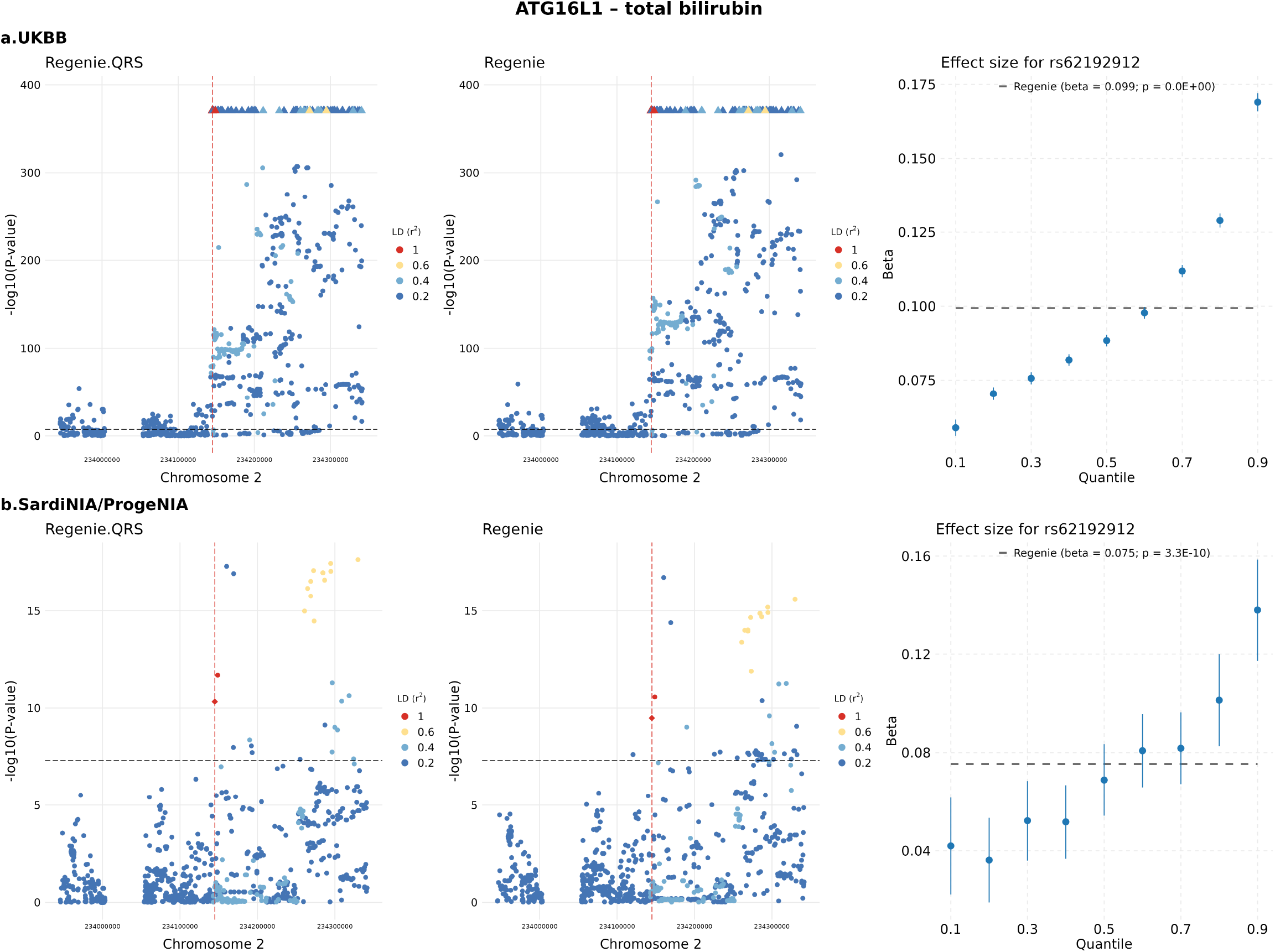
Heterogeneous effect of rs62192912 located at the *ATG16L1* locus and total bilirubin. Locus zoom plots display − log_10_(p-value) for each variant, with the lead SNP at each locus shown as a diamond and highlighted by a vertical dashed line. Colors indicate the linkage disequilibrium (LD, *r*^2^) between each variant and the lead SNP. **a**. UKBB; **b**. Sardinia. The horizontal dashed line marks the genome-wide significance threshold at 5 *×* 10^−8^. For the lead Regenie.QRS SNP, the quantile-specific effect sizes *±* standard errors are shown in the right panel at different quantile levels.

**Figure S5:**
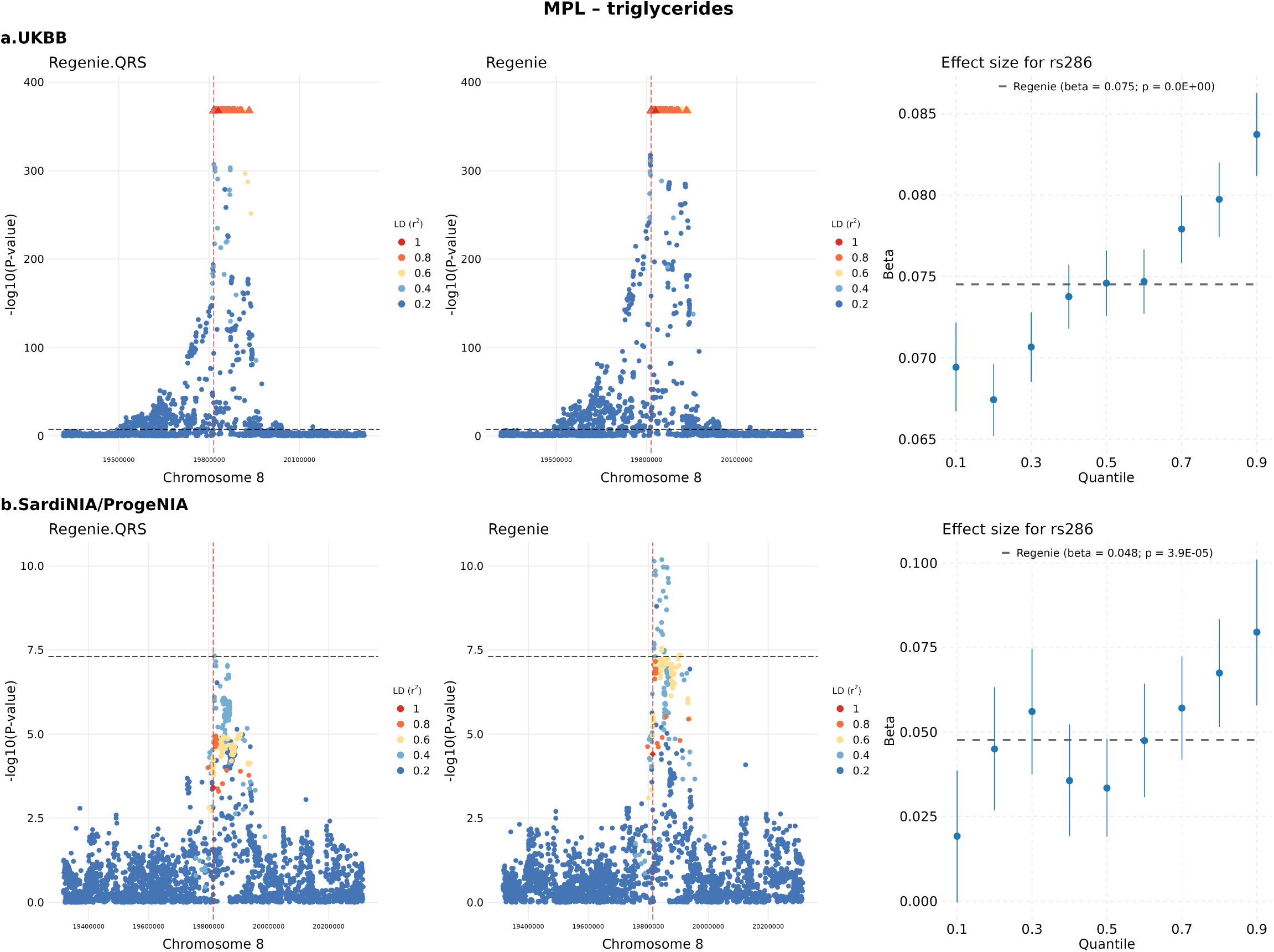
Heterogeneous effect of rs286 located at the *MPL* locus and triglycerides. Locus zoom plots display −log_10_(p-value) for each variant, with the lead SNP at each locus shown as a diamond and highlighted by a vertical dashed line. Colors indicate the linkage disequilibrium (LD, *r*^2^) between each variant and the lead SNP. **a**. UKBB; **b**. Sardinia. The horizontal dashed line marks the genome-wide significance threshold at 5 *×* 10^−8^. For the lead Regenie.QRS SNP, the quantile-specific effect sizes *±* standard errors are shown in the right panel at different quantile levels.

**Table S1:** Empirical type I error rates with 409,627 white British individuals under the homogeneous model with normal error distribution.

| Quantiles | | $10^{-8}$ | $10^{-7}$ | $10^{-6}$ | $10^{-5}$ | $10^{-4}$ | 0.001 | 0.005 | 0.01 | 0.05 |
| --- | --- | --- | --- | --- | --- | --- | --- | --- | --- | --- |
| | | $(0, 5.74 \times 10^{-8})$ | $(0, 2.50 \times 10^{-7})$ | $(5.26 \times 10^{-7}, 1.47 \times 10^{-6})$ | $(8.50 \times 10^{-6}, 1.15 \times 10^{-5})$ | $(9.53 \times 10^{-5}, 1.05 \times 10^{-4})$ | $(0.0000985, 0.00101)$ | $(0.00497, 0.00503)$ | $(0.00995, 0.01005)$ | $(0.0499, 0.0501)$ |
| Regenie | - | $1.22 \times 10^{-7}$ | $2.44 \times 10^{-7}$ | $1.46 \times 10^{-6}$ | $1.35 \times 10^{-5}$ | $1.16 \times 10^{-4}$ | 0.0011 | 0.0052 | 0.0104 | 0.0510 |
| QRS | composite | $2.38 \times 10^{-6}$ | $3.53 \times 10^{-6}$ | $5.48 \times 10^{-6}$ | $1.86 \times 10^{-5}$ | $1.28 \times 10^{-4}$ | 0.0012 | 0.0062 | 0.0124 | 0.0594 |
| | 0.2 | $1.22 \times 10^{-6}$ | $1.83 \times 10^{-6}$ | $3.41 \times 10^{-6}$ | $1.61 \times 10^{-5}$ | $1.19 \times 10^{-4}$ | 0.0011 | 0.0054 | 0.0107 | 0.0520 |
| | 0.5 | $1.89 \times 10^{-6}$ | $2.74 \times 10^{-6}$ | $5.48 \times 10^{-6}$ | $1.65 \times 10^{-5}$ | $1.24 \times 10^{-4}$ | 0.0011 | 0.0054 | 0.0107 | 0.0522 |
| | 0.8 | $1.40 \times 10^{-6}$ | $2.13 \times 10^{-6}$ | $3.84 \times 10^{-6}$ | $1.57 \times 10^{-5}$ | $1.19 \times 10^{-4}$ | 0.0011 | 0.0054 | 0.0107 | 0.0519 |
| Regenie.QRS | composite | $< 5.84 \times 10^{-8}$ | $6.09 \times 10^{-8}$ | $1.34 \times 10^{-6}$ | $1.35 \times 10^{-5}$ | $1.16 \times 10^{-4}$ | 0.0011 | 0.0057 | 0.0116 | 0.0569 |
| | 0.2 | $< 5.84 \times 10^{-8}$ | $< 5.84 \times 10^{-8}$ | $9.74 \times 10^{-7}$ | $1.12 \times 10^{-5}$ | $1.05 \times 10^{-4}$ | 0.0010 | 0.0051 | 0.0101 | 0.0504 |
| | 0.5 | $< 5.84 \times 10^{-8}$ | $4.26 \times 10^{-7}$ | $1.58 \times 10^{-6}$ | $1.21 \times 10^{-5}$ | $1.04 \times 10^{-4}$ | 0.0010 | 0.0051 | 0.0102 | 0.0507 |
| | 0.8 | $< 5.84 \times 10^{-8}$ | $< 5.84 \times 10^{-8}$ | $9.74 \times 10^{-7}$ | $1.11 \times 10^{-5}$ | $1.07 \times 10^{-4}$ | 0.0010 | 0.0051 | 0.0102 | 0.0505 |
| BLUP.QRS | composite | $1.40 \times 10^{-6}$ | $1.71 \times 10^{-6}$ | $3.05 \times 10^{-6}$ | $1.58 \times 10^{-5}$ | $1.24 \times 10^{-4}$ | 0.0012 | 0.0060 | 0.0121 | 0.0585 |
| | 0.2 | $7.31 \times 10^{-7}$ | $1.28 \times 10^{-6}$ | $2.80 \times 10^{-6}$ | $1.41 \times 10^{-5}$ | $1.14 \times 10^{-4}$ | 0.0011 | 0.0053 | 0.0105 | 0.0514 |
| | 0.5 | $1.22 \times 10^{-6}$ | $1.46 \times 10^{-6}$ | $3.41 \times 10^{-6}$ | $1.38 \times 10^{-5}$ | $1.16 \times 10^{-4}$ | 0.0011 | 0.0053 | 0.0105 | 0.0516 |
| | 0.8 | $5.48 \times 10^{-7}$ | $7.92 \times 10^{-7}$ | $2.01 \times 10^{-6}$ | $1.35 \times 10^{-5}$ | $1.12 \times 10^{-4}$ | 0.0011 | 0.0053 | 0.0105 | 0.0515 |

**Table S2:** Empirical type I error rates with 409,627 white British individuals under the *t*_(2)_ error model.

| Quantiles | | $10^{-8}$ | $10^{-7}$ | $10^{-6}$ | $10^{-5}$ | $10^{-4}$ | 0.001 | 0.005 | 0.01 | 0.05 |
| --- | --- | --- | --- | --- | --- | --- | --- | --- | --- | --- |
| | | $(0, 5.74 \times 10^{-8})$ | $(0, 2.50 \times 10^{-7})$ | $(5.26 \times 10^{-7}, 1.47 \times 10^{-6})$ | $(8.50 \times 10^{-6}, 1.15 \times 10^{-5})$ | $(9.53 \times 10^{-5}, 1.05 \times 10^{-4})$ | $(0.0000985, 0.00101)$ | $(0.00497, 0.00503)$ | $(0.00995, 0.01005)$ | $(0.0499, 0.0501)$ |
| Regenie | - | $7.92 \times 10^{-7}$ | $1.22 \times 10^{-6}$ | $2.92 \times 10^{-6}$ | $1.62 \times 10^{-5}$ | $1.21 \times 10^{-4}$ | 0.0011 | 0.0053 | 0.0105 | 0.0514 |
| QRS | composite | $7.13 \times 10^{-6}$ | $9.26 \times 10^{-6}$ | $1.49 \times 10^{-5}$ | $3.47 \times 10^{-5}$ | $1.63 \times 10^{-4}$ | 0.0013 | 0.0064 | 0.0126 | 0.0599 |
| | 0.2 | $3.29 \times 10^{-6}$ | $5.91 \times 10^{-6}$ | $1.22 \times 10^{-5}$ | $2.87 \times 10^{-5}$ | $1.48 \times 10^{-4}$ | 0.0012 | 0.0055 | 0.0108 | 0.0524 |
| | 0.5 | $5.91 \times 10^{-6}$ | $9.14 \times 10^{-6}$ | $1.37 \times 10^{-5}$ | $3.10 \times 10^{-5}$ | $1.50 \times 10^{-4}$ | 0.0012 | 0.0056 | 0.0109 | 0.0528 |
| | 0.8 | $4.14 \times 10^{-6}$ | $4.87 \times 10^{-6}$ | $7.73 \times 10^{-6}$ | $2.24 \times 10^{-5}$ | $1.32 \times 10^{-4}$ | 0.0012 | 0.0055 | 0.0108 | 0.0523 |
| Regenie.QRS | composite | $1.16 \times 10^{-6}$ | $1.64 \times 10^{-6}$ | $3.53 \times 10^{-6}$ | $1.51 \times 10^{-5}$ | $1.25 \times 10^{-4}$ | 0.0012 | 0.0059 | 0.0119 | 0.0579 |
| | 0.2 | $7.31 \times 10^{-7}$ | $9.14 \times 10^{-7}$ | $1.95 \times 10^{-6}$ | $1.13 \times 10^{-5}$ | $1.08 \times 10^{-4}$ | 0.0011 | 0.0052 | 0.0104 | 0.0511 |
| | 0.5 | $1.10 \times 10^{-6}$ | $1.58 \times 10^{-6}$ | $3.23 \times 10^{-6}$ | $1.75 \times 10^{-5}$ | $1.27 \times 10^{-4}$ | 0.0011 | 0.0053 | 0.0105 | 0.0515 |
| | 0.8 | $3.05 \times 10^{-7}$ | $7.31 \times 10^{-7}$ | $1.71 \times 10^{-6}$ | $1.32 \times 10^{-5}$ | $1.14 \times 10^{-4}$ | 0.0011 | 0.0052 | 0.0103 | 0.0509 |
| BLUP.QRS | composite | $2.25 \times 10^{-6}$ | $3.29 \times 10^{-6}$ | $5.24 \times 10^{-6}$ | $1.71 \times 10^{-5}$ | $1.28 \times 10^{-4}$ | 0.0012 | 0.0061 | 0.0122 | 0.0591 |
| | 0.2 | $1.77 \times 10^{-6}$ | $2.01 \times 10^{-6}$ | $3.53 \times 10^{-6}$ | $1.57 \times 10^{-5}$ | $1.26 \times 10^{-4}$ | 0.0011 | 0.0054 | 0.0106 | 0.0518 |
| | 0.5 | $2.13 \times 10^{-6}$ | $2.50 \times 10^{-6}$ | $4.38 \times 10^{-6}$ | $1.67 \times 10^{-5}$ | $1.25 \times 10^{-4}$ | 0.0011 | 0.0054 | 0.0107 | 0.0520 |
| | 0.8 | $1.34 \times 10^{-6}$ | $2.01 \times 10^{-6}$ | $3.84 \times 10^{-6}$ | $1.52 \times 10^{-5}$ | $1.21 \times 10^{-4}$ | 0.0011 | 0.0054 | 0.0106 | 0.0518 |

**Table S3:**
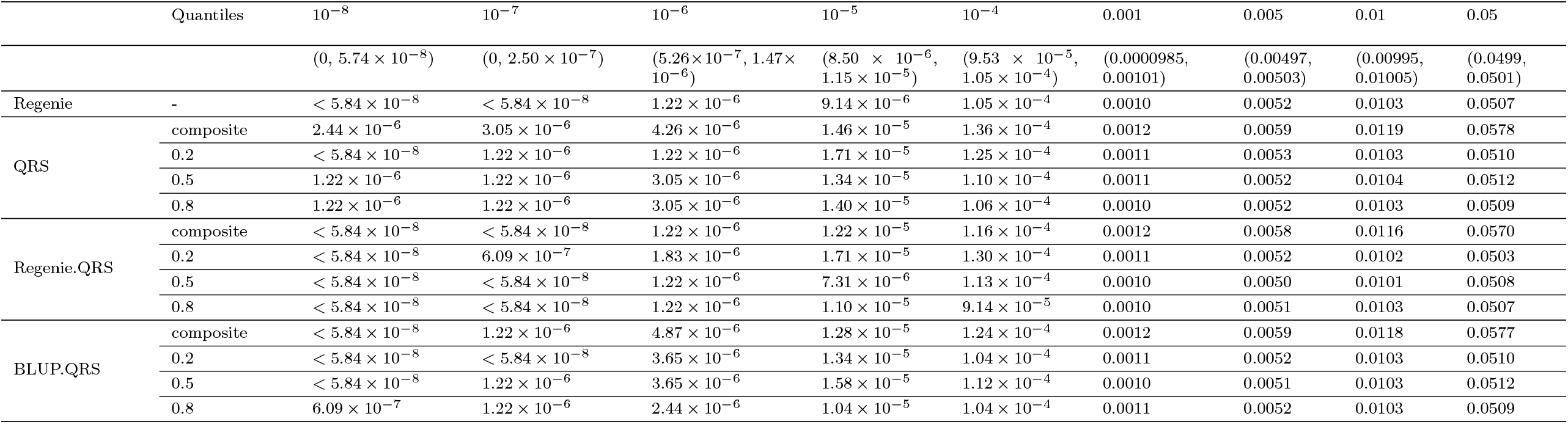
Empirical type I error rates with 409,627 white British individuals under the dominance model.

| Quantiles | | $10^{-8}$ | $10^{-7}$ | $10^{-6}$ | $10^{-5}$ | $10^{-4}$ | 0.001 | 0.005 | 0.01 | 0.05 |
| --- | --- | --- | --- | --- | --- | --- | --- | --- | --- | --- |
| | | $(0, 5.74 \times 10^{-8})$ | $(0, 2.50 \times 10^{-7})$ | $(5.26 \times 10^{-7}, 1.47 \times 10^{-6})$ | $(8.50 \times 10^{-6}, 1.15 \times 10^{-5})$ | $(9.53 \times 10^{-5}, 1.05 \times 10^{-4})$ | $(0.0000985, 0.00101)$ | $(0.00497, 0.00503)$ | $(0.00995, 0.01005)$ | $(0.0499, 0.0501)$ |
| Regenie | - | $< 5.84 \times 10^{-8}$ | $< 5.84 \times 10^{-8}$ | $1.22 \times 10^{-6}$ | $9.14 \times 10^{-6}$ | $1.05 \times 10^{-4}$ | 0.0010 | 0.0052 | 0.0103 | 0.0507 |
| QRS | composite | $2.44 \times 10^{-6}$ | $3.05 \times 10^{-6}$ | $4.26 \times 10^{-6}$ | $1.46 \times 10^{-5}$ | $1.36 \times 10^{-4}$ | 0.0012 | 0.0059 | 0.0119 | 0.0578 |
| | 0.2 | $< 5.84 \times 10^{-8}$ | $1.22 \times 10^{-6}$ | $1.22 \times 10^{-6}$ | $1.71 \times 10^{-5}$ | $1.25 \times 10^{-4}$ | 0.0011 | 0.0053 | 0.0103 | 0.0510 |
| | 0.5 | $1.22 \times 10^{-6}$ | $1.22 \times 10^{-6}$ | $3.05 \times 10^{-6}$ | $1.34 \times 10^{-5}$ | $1.10 \times 10^{-4}$ | 0.0011 | 0.0052 | 0.0104 | 0.0512 |
| | 0.8 | $1.22 \times 10^{-6}$ | $1.22 \times 10^{-6}$ | $3.05 \times 10^{-6}$ | $1.40 \times 10^{-5}$ | $1.06 \times 10^{-4}$ | 0.0010 | 0.0052 | 0.0103 | 0.0509 |
| Regenie.QRS | composite | $< 5.84 \times 10^{-8}$ | $< 5.84 \times 10^{-8}$ | $1.22 \times 10^{-6}$ | $1.22 \times 10^{-5}$ | $1.16 \times 10^{-4}$ | 0.0012 | 0.0058 | 0.0116 | 0.0570 |
| | 0.2 | $< 5.84 \times 10^{-8}$ | $6.09 \times 10^{-7}$ | $1.83 \times 10^{-6}$ | $1.71 \times 10^{-5}$ | $1.30 \times 10^{-4}$ | 0.0011 | 0.0052 | 0.0102 | 0.0503 |
| | 0.5 | $< 5.84 \times 10^{-8}$ | $< 5.84 \times 10^{-8}$ | $1.22 \times 10^{-6}$ | $7.31 \times 10^{-6}$ | $1.13 \times 10^{-4}$ | 0.0010 | 0.0050 | 0.0101 | 0.0508 |
| | 0.8 | $< 5.84 \times 10^{-8}$ | $< 5.84 \times 10^{-8}$ | $1.22 \times 10^{-6}$ | $1.10 \times 10^{-5}$ | $9.14 \times 10^{-5}$ | 0.0010 | 0.0051 | 0.0103 | 0.0507 |
| BLUP.QRS | composite | $< 5.84 \times 10^{-8}$ | $1.22 \times 10^{-6}$ | $4.87 \times 10^{-6}$ | $1.28 \times 10^{-5}$ | $1.24 \times 10^{-4}$ | 0.0012 | 0.0059 | 0.0118 | 0.0577 |
| | 0.2 | $< 5.84 \times 10^{-8}$ | $< 5.84 \times 10^{-8}$ | $3.65 \times 10^{-6}$ | $1.34 \times 10^{-5}$ | $1.04 \times 10^{-4}$ | 0.0011 | 0.0052 | 0.0103 | 0.0510 |
| | 0.5 | $< 5.84 \times 10^{-8}$ | $1.22 \times 10^{-6}$ | $3.65 \times 10^{-6}$ | $1.58 \times 10^{-5}$ | $1.12 \times 10^{-4}$ | 0.0010 | 0.0051 | 0.0103 | 0.0512 |
| | 0.8 | $6.09 \times 10^{-7}$ | $1.22 \times 10^{-6}$ | $2.44 \times 10^{-6}$ | $1.04 \times 10^{-5}$ | $1.04 \times 10^{-4}$ | 0.0011 | 0.0052 | 0.0103 | 0.0509 |

**Table S4:** Empirical type I error rates with 409,627 white British individuals under the heterogeneous model.

| Quantiles | | $10^{-8}$ | $10^{-7}$ | $10^{-6}$ | $10^{-5}$ | $10^{-4}$ | 0.001 | 0.005 | 0.01 | 0.05 |
| --- | --- | --- | --- | --- | --- | --- | --- | --- | --- | --- |
| | | $(0, 5.74 \times 10^{-8})$ | $(0, 2.50 \times 10^{-7})$ | $(5.26 \times 10^{-7}, 1.47 \times 10^{-6})$ | $(8.50 \times 10^{-6}, 1.15 \times 10^{-5})$ | $(9.53 \times 10^{-5}, 1.05 \times 10^{-4})$ | $(0.0000985, 0.00101)$ | $(0.00497, 0.00503)$ | $(0.00995, 0.01005)$ | $(0.0499, 0.0501)$ |
| Regenie | - | $6.09 \times 10^{-8}$ | $3.05 \times 10^{-7}$ | $2.01 \times 10^{-6}$ | $1.35 \times 10^{-5}$ | $1.04 \times 10^{-4}$ | 0.0010 | 0.0050 | 0.0100 | 0.0499 |
| QRS | composite | $1.28 \times 10^{-6}$ | $1.83 \times 10^{-6}$ | $3.72 \times 10^{-6}$ | $1.55 \times 10^{-5}$ | $1.22 \times 10^{-4}$ | 0.0012 | 0.0059 | 0.0119 | 0.0580 |
| | 0.2 | $4.26 \times 10^{-7}$ | $7.31 \times 10^{-7}$ | $2.25 \times 10^{-6}$ | $1.18 \times 10^{-5}$ | $1.06 \times 10^{-4}$ | 0.0011 | 0.0052 | 0.0104 | 0.0511 |
| | 0.5 | $1.10 \times 10^{-6}$ | $1.58 \times 10^{-6}$ | $2.98 \times 10^{-6}$ | $1.39 \times 10^{-5}$ | $1.13 \times 10^{-4}$ | 0.0011 | 0.0053 | 0.0105 | 0.0514 |
| | 0.8 | $5.48 \times 10^{-7}$ | $9.14 \times 10^{-7}$ | $2.25 \times 10^{-6}$ | $1.29 \times 10^{-5}$ | $1.09 \times 10^{-4}$ | 0.0011 | 0.0052 | 0.0104 | 0.0511 |
| Regenie.QRS | composite | $6.09 \times 10^{-8}$ | $1.22 \times 10^{-7}$ | $9.14 \times 10^{-7}$ | $1.02 \times 10^{-5}$ | $1.05 \times 10^{-4}$ | 0.0011 | 0.0056 | 0.0113 | 0.0560 |
| | 0.2 | $< 5.84 \times 10^{-8}$ | $6.09 \times 10^{-8}$ | $1.16 \times 10^{-6}$ | $9.26 \times 10^{-6}$ | $9.80 \times 10^{-5}$ | 0.0010 | 0.0050 | 0.0100 | 0.0499 |
| | 0.5 | $6.09 \times 10^{-8}$ | $2.44 \times 10^{-7}$ | $1.28 \times 10^{-6}$ | $1.07 \times 10^{-5}$ | $9.91 \times 10^{-5}$ | 0.0010 | 0.0050 | 0.0100 | 0.0501 |
| | 0.8 | $< 5.84 \times 10^{-8}$ | $< 5.84 \times 10^{-8}$ | $1.10 \times 10^{-6}$ | $1.07 \times 10^{-5}$ | $1.03 \times 10^{-4}$ | 0.0010 | 0.0050 | 0.0100 | 0.0500 |
| BLUP.QRS | composite | $6.70 \times 10^{-7}$ | $1.22 \times 10^{-6}$ | $2.86 \times 10^{-6}$ | $1.45 \times 10^{-5}$ | $1.16 \times 10^{-4}$ | 0.0012 | 0.0058 | 0.0118 | 0.0573 |
| | 0.2 | $2.44 \times 10^{-7}$ | $5.48 \times 10^{-7}$ | $2.01 \times 10^{-6}$ | $1.19 \times 10^{-5}$ | $1.11 \times 10^{-4}$ | 0.0011 | 0.0052 | 0.0103 | 0.0507 |
| | 0.5 | $6.09 \times 10^{-7}$ | $1.34 \times 10^{-6}$ | $2.56 \times 10^{-6}$ | $1.49 \times 10^{-5}$ | $1.13 \times 10^{-4}$ | 0.0011 | 0.0052 | 0.0103 | 0.0510 |
| | 0.8 | $1.83 \times 10^{-7}$ | $4.87 \times 10^{-7}$ | $1.95 \times 10^{-6}$ | $1.13 \times 10^{-5}$ | $1.07 \times 10^{-4}$ | 0.0010 | 0.0052 | 0.0103 | 0.0507 |

**Table S5:** Empirical type I error rates with 46,887 first-degree white British relatives under the homogeneous model with normal error distribution.

| Quantiles |  | 1e-08 | 1e-07 | 1e-06 | 1e-05 | 1e-04 | 0.001 | 0.005 | 0.01 | 0.05 |
| --- | --- | --- | --- | --- | --- | --- | --- | --- | --- | --- |
| | | $(0, 5.74 \times 10^{-8})$ | $(0, 2.50 \times 10^{-7})$ | $(5.26 \times 10^{-7}, 1.47 \times 10^{-6})$ | $(8.50 \times 10^{-6}, 1.15 \times 10^{-5})$ | $(9.53 \times 10^{-5}, 1.05 \times 10^{-4})$ | $(0.0000985, 0.00101)$ | $(0.00497, 0.00503)$ | $(0.00995, 0.01005)$ | $(0.0499, 0.0501)$ |
| Regenie | - | $< 5.84 \times 10^{-8}$ | $1.17 \times 10^{-7}$ | $6.43 \times 10^{-7}$ | $7.01 \times 10^{-6}$ | $7.29 \times 10^{-5}$ | 0.0008 | 0.0044 | 0.0090 | 0.0468 |
| QRS | composite | $< 5.84 \times 10^{-8}$ | $2.34 \times 10^{-7}$ | $3.27 \times 10^{-6}$ | $3.39 \times 10^{-5}$ | $2.78 \times 10^{-4}$ | 0.0022 | 0.0098 | 0.0184 | 0.0768 |
| | 0.2 | $< 5.84 \times 10^{-8}$ | $2.92 \times 10^{-7}$ | $3.15 \times 10^{-6}$ | $2.72 \times 10^{-5}$ | $2.12 \times 10^{-4}$ | 0.0017 | 0.0076 | 0.0143 | 0.0623 |
| | 0.5 | $5.84 \times 10^{-8}$ | $2.34 \times 10^{-7}$ | $3.51 \times 10^{-6}$ | $3.09 \times 10^{-5}$ | $2.45 \times 10^{-4}$ | 0.0019 | 0.0081 | 0.0151 | 0.0643 |
| | 0.8 | $< 5.84 \times 10^{-8}$ | $3.51 \times 10^{-7}$ | $2.86 \times 10^{-6}$ | $2.52 \times 10^{-5}$ | $2.13 \times 10^{-4}$ | 0.0018 | 0.0076 | 0.0142 | 0.0621 |
| Regenie.QRS | composite | $< 5.84 \times 10^{-8}$ | $5.84 \times 10^{-8}$ | $5.26 \times 10^{-7}$ | $7.48 \times 10^{-6}$ | $8.92 \times 10^{-5}$ | 0.0009 | 0.0050 | 0.0103 | 0.0530 |
| | 0.2 | $< 5.84 \times 10^{-8}$ | $< 5.84 \times 10^{-8}$ | $1.05 \times 10^{-6}$ | $8.35 \times 10^{-6}$ | $8.95 \times 10^{-5}$ | 0.0009 | 0.0047 | 0.0095 | 0.0484 |
| | 0.5 | $< 5.84 \times 10^{-8}$ | $< 5.84 \times 10^{-8}$ | $8.76 \times 10^{-7}$ | $8.00 \times 10^{-6}$ | $7.78 \times 10^{-5}$ | 0.0009 | 0.0045 | 0.0091 | 0.0471 |
| | 0.8 | $< 5.84 \times 10^{-8}$ | $< 5.84 \times 10^{-8}$ | $3.51 \times 10^{-7}$ | $7.89 \times 10^{-6}$ | $9.14 \times 10^{-5}$ | 0.0009 | 0.0047 | 0.0095 | 0.0483 |
| BLUP.QRS | composite | $< 5.84 \times 10^{-8}$ | $2.92 \times 10^{-7}$ | $2.10 \times 10^{-6}$ | $2.09 \times 10^{-5}$ | $1.90 \times 10^{-4}$ | 0.0017 | 0.0079 | 0.0153 | 0.0684 |
| | 0.2 | $1.75 \times 10^{-7}$ | $3.51 \times 10^{-7}$ | $2.40 \times 10^{-6}$ | $1.96 \times 10^{-5}$ | $1.67 \times 10^{-4}$ | 0.0014 | 0.0065 | 0.0125 | 0.0572 |
| | 0.5 | $< 5.84 \times 10^{-8}$ | $5.84 \times 10^{-8}$ | $2.63 \times 10^{-6}$ | $1.92 \times 10^{-5}$ | $1.74 \times 10^{-4}$ | 0.0015 | 0.0068 | 0.0130 | 0.0587 |
| | 0.8 | $< 5.84 \times 10^{-8}$ | $2.34 \times 10^{-7}$ | $2.04 \times 10^{-6}$ | $1.80 \times 10^{-5}$ | $1.59 \times 10^{-4}$ | 0.0014 | 0.0064 | 0.0124 | 0.0571 |

**Table S6:**
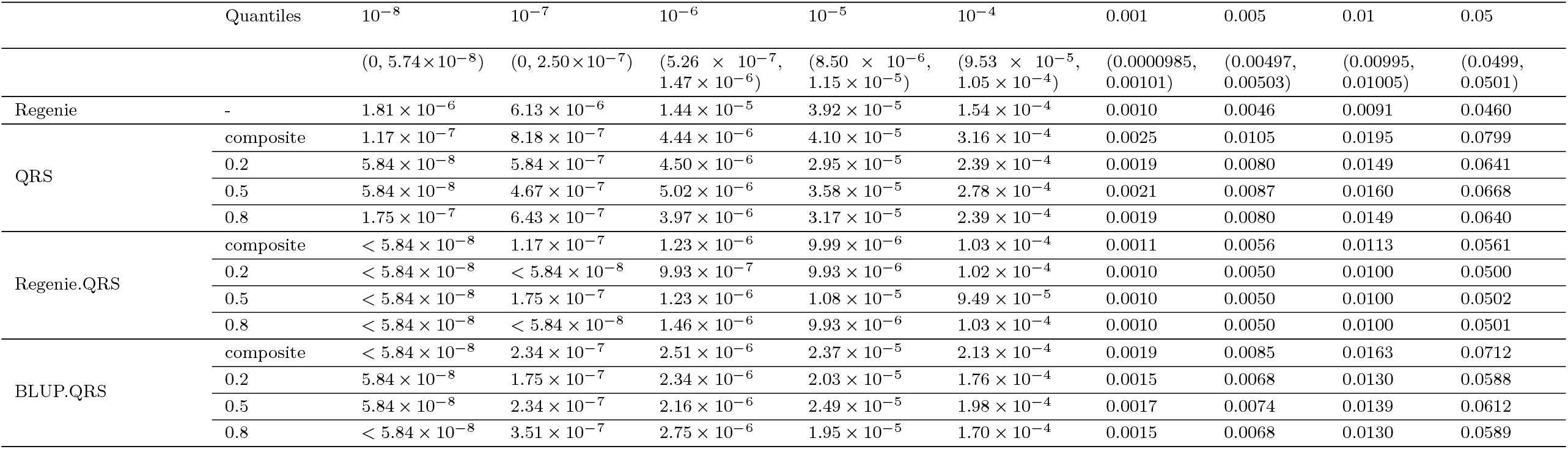
Empirical type I error rates with 46,887 first-degree white British relatives under the *t*_(2)_ error model.

**Table S7:** Empirical type I error rates with 46,887 first-degree white British relatives under the dominance model.

| Quantiles | | $10^{-8}$ | $10^{-7}$ | $10^{-6}$ | $10^{-5}$ | $10^{-4}$ | 0.001 | 0.005 | 0.01 | 0.05 |
| --- | --- | --- | --- | --- | --- | --- | --- | --- | --- | --- |
| | | $(0, 5.74 \times 10^{-8})$ | $(0, 2.50 \times 10^{-7})$ | $(5.26 \times 10^{-7}, 1.47 \times 10^{-6})$ | $(8.50 \times 10^{-6}, 1.15 \times 10^{-5})$ | $(9.53 \times 10^{-5}, 1.05 \times 10^{-4})$ | $(0.0000985, 0.00101)$ | $(0.00497, 0.00503)$ | $(0.00995, 0.01005)$ | $(0.0499, 0.0501)$ |
| Regenie | - | $< 5.84 \times 10^{-8}$ | $1.17 \times 10^{-7}$ | $1.40 \times 10^{-6}$ | $9.52 \times 10^{-6}$ | $9.06 \times 10^{-5}$ | 0.0009 | 0.0048 | 0.0096 | 0.0488 |
| QRS | composite | $< 5.84 \times 10^{-8}$ | $6.43 \times 10^{-7}$ | $4.50 \times 10^{-6}$ | $3.45 \times 10^{-5}$ | $2.49 \times 10^{-4}$ | 0.0021 | 0.0091 | 0.0173 | 0.0742 |
| | 0.2 | $< 5.84 \times 10^{-8}$ | $1.75 \times 10^{-7}$ | $3.10 \times 10^{-6}$ | $2.32 \times 10^{-5}$ | $1.96 \times 10^{-4}$ | 0.0016 | 0.0072 | 0.0136 | 0.0605 |
| | 0.5 | $1.75 \times 10^{-7}$ | $6.43 \times 10^{-7}$ | $4.21 \times 10^{-6}$ | $2.55 \times 10^{-5}$ | $2.16 \times 10^{-4}$ | 0.0017 | 0.0076 | 0.0143 | 0.0623 |
| | 0.8 | $5.84 \times 10^{-8}$ | $4.09 \times 10^{-7}$ | $3.04 \times 10^{-6}$ | $2.45 \times 10^{-5}$ | $1.97 \times 10^{-4}$ | 0.0017 | 0.0072 | 0.0137 | 0.0609 |
| Regenie.QRS | composite | $< 5.84 \times 10^{-8}$ | $1.17 \times 10^{-7}$ | $1.23 \times 10^{-6}$ | $1.07 \times 10^{-5}$ | $1.06 \times 10^{-4}$ | 0.0011 | 0.0057 | 0.0114 | 0.0566 |
| | 0.2 | $< 5.84 \times 10^{-8}$ | $1.75 \times 10^{-7}$ | $9.35 \times 10^{-7}$ | $1.10 \times 10^{-5}$ | $1.07 \times 10^{-4}$ | 0.0010 | 0.0051 | 0.0101 | 0.0503 |
| | 0.5 | $< 5.84 \times 10^{-8}$ | $5.84 \times 10^{-8}$ | $8.18 \times 10^{-7}$ | $9.93 \times 10^{-6}$ | $9.23 \times 10^{-5}$ | 0.0010 | 0.0049 | 0.0099 | 0.0498 |
| | 0.8 | $< 5.84 \times 10^{-8}$ | $5.84 \times 10^{-8}$ | $8.76 \times 10^{-7}$ | $9.58 \times 10^{-6}$ | $1.07 \times 10^{-4}$ | 0.0010 | 0.0051 | 0.0102 | 0.0506 |
| BLUP.QRS | composite | $5.84 \times 10^{-8}$ | $1.17 \times 10^{-7}$ | $2.75 \times 10^{-6}$ | $2.33 \times 10^{-5}$ | $1.89 \times 10^{-4}$ | 0.0017 | 0.0079 | 0.0153 | 0.0686 |
| | 0.2 | $5.84 \times 10^{-8}$ | $1.17 \times 10^{-7}$ | $1.99 \times 10^{-6}$ | $1.90 \times 10^{-5}$ | $1.66 \times 10^{-4}$ | 0.0014 | 0.0065 | 0.0124 | 0.0572 |
| | 0.5 | $< 5.84 \times 10^{-8}$ | $1.75 \times 10^{-7}$ | $2.51 \times 10^{-6}$ | $2.09 \times 10^{-5}$ | $1.69 \times 10^{-4}$ | 0.0015 | 0.0067 | 0.0129 | 0.0586 |
| | 0.8 | $< 5.84 \times 10^{-8}$ | $2.34 \times 10^{-7}$ | $2.04 \times 10^{-6}$ | $1.72 \times 10^{-5}$ | $1.64 \times 10^{-4}$ | 0.0014 | 0.0065 | 0.0125 | 0.0575 |

**Table S8:**
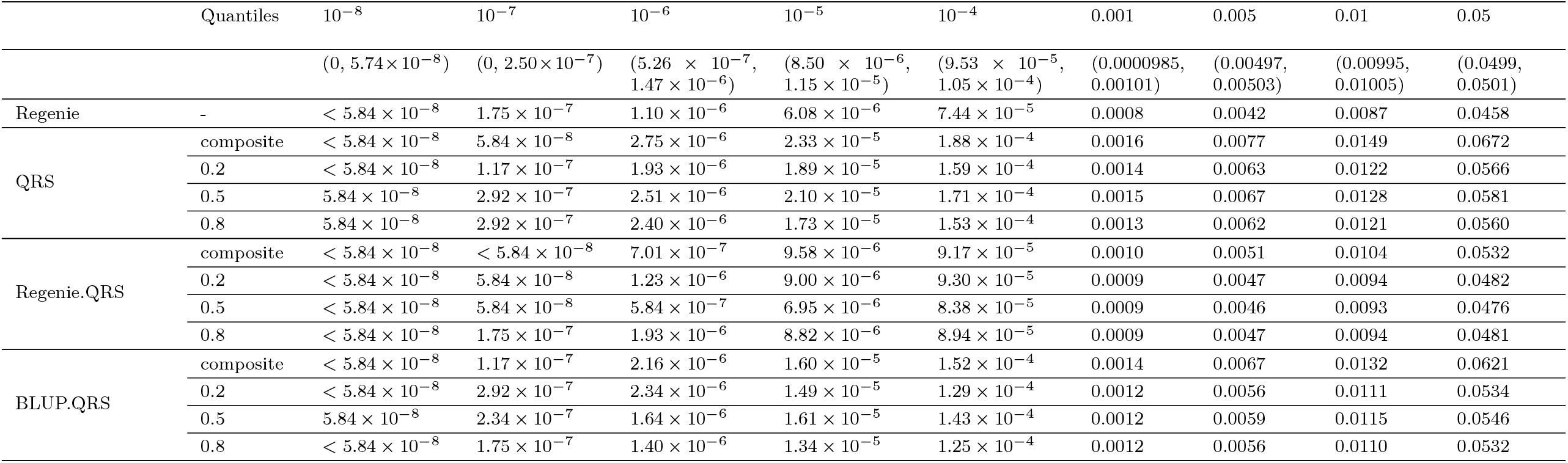
Empirical type I error rates with 46,887 first-degree white British relatives under the heterogeneous model.

| | Quantiles | $10^{-8}$ | $10^{-7}$ | $10^{-6}$ | $10^{-5}$ | $10^{-4}$ | 0.001 | 0.005 | 0.01 | 0.05 |
| --- | --- | --- | --- | --- | --- | --- | --- | --- | --- | --- |
| | | $(0, 5.74 \times 10^{-8})$ | $(0, 2.50 \times 10^{-7})$ | $(5.26 \times 10^{-7}, 1.47 \times 10^{-6})$ | $(8.50 \times 10^{-6}, 1.15 \times 10^{-5})$ | $(9.53 \times 10^{-5}, 1.05 \times 10^{-4})$ | $(0.0000985, 0.00101)$ | $(0.00497, 0.00503)$ | $(0.00995, 0.01005)$ | $(0.0499, 0.0501)$ |
| Regenie | - | $< 5.84 \times 10^{-8}$ | $1.75 \times 10^{-7}$ | $1.10 \times 10^{-6}$ | $6.08 \times 10^{-6}$ | $7.44 \times 10^{-5}$ | 0.0008 | 0.0042 | 0.0087 | 0.0458 |
| QRS | composite | $< 5.84 \times 10^{-8}$ | $5.84 \times 10^{-8}$ | $2.75 \times 10^{-6}$ | $2.33 \times 10^{-5}$ | $1.88 \times 10^{-4}$ | 0.0016 | 0.0077 | 0.0149 | 0.0672 |
| | 0.2 | $< 5.84 \times 10^{-8}$ | $1.17 \times 10^{-7}$ | $1.93 \times 10^{-6}$ | $1.89 \times 10^{-5}$ | $1.59 \times 10^{-4}$ | 0.0014 | 0.0063 | 0.0122 | 0.0566 |
| | 0.5 | $5.84 \times 10^{-8}$ | $2.92 \times 10^{-7}$ | $2.51 \times 10^{-6}$ | $2.10 \times 10^{-5}$ | $1.71 \times 10^{-4}$ | 0.0015 | 0.0067 | 0.0128 | 0.0581 |
| | 0.8 | $5.84 \times 10^{-8}$ | $2.92 \times 10^{-7}$ | $2.40 \times 10^{-6}$ | $1.73 \times 10^{-5}$ | $1.53 \times 10^{-4}$ | 0.0013 | 0.0062 | 0.0121 | 0.0560 |
| Regenie.QRS | composite | $< 5.84 \times 10^{-8}$ | $< 5.84 \times 10^{-8}$ | $7.01 \times 10^{-7}$ | $9.58 \times 10^{-6}$ | $9.17 \times 10^{-5}$ | 0.0010 | 0.0051 | 0.0104 | 0.0532 |
| | 0.2 | $< 5.84 \times 10^{-8}$ | $5.84 \times 10^{-8}$ | $1.23 \times 10^{-6}$ | $9.00 \times 10^{-6}$ | $9.30 \times 10^{-5}$ | 0.0009 | 0.0047 | 0.0094 | 0.0482 |
| | 0.5 | $< 5.84 \times 10^{-8}$ | $5.84 \times 10^{-8}$ | $5.84 \times 10^{-7}$ | $6.95 \times 10^{-6}$ | $8.38 \times 10^{-5}$ | 0.0009 | 0.0046 | 0.0093 | 0.0476 |
| | 0.8 | $< 5.84 \times 10^{-8}$ | $1.75 \times 10^{-7}$ | $1.93 \times 10^{-6}$ | $8.82 \times 10^{-6}$ | $8.94 \times 10^{-5}$ | 0.0009 | 0.0047 | 0.0094 | 0.0481 |
| BLUP.QRS | composite | $< 5.84 \times 10^{-8}$ | $1.17 \times 10^{-7}$ | $2.16 \times 10^{-6}$ | $1.60 \times 10^{-5}$ | $1.52 \times 10^{-4}$ | 0.0014 | 0.0067 | 0.0132 | 0.0621 |
| | 0.2 | $< 5.84 \times 10^{-8}$ | $2.92 \times 10^{-7}$ | $2.34 \times 10^{-6}$ | $1.49 \times 10^{-5}$ | $1.29 \times 10^{-4}$ | 0.0012 | 0.0056 | 0.0111 | 0.0534 |
| | 0.5 | $5.84 \times 10^{-8}$ | $2.34 \times 10^{-7}$ | $1.64 \times 10^{-6}$ | $1.61 \times 10^{-5}$ | $1.43 \times 10^{-4}$ | 0.0012 | 0.0059 | 0.0115 | 0.0546 |
| | 0.8 | $< 5.84 \times 10^{-8}$ | $1.75 \times 10^{-7}$ | $1.40 \times 10^{-6}$ | $1.34 \times 10^{-5}$ | $1.25 \times 10^{-4}$ | 0.0012 | 0.0056 | 0.0110 | 0.0532 |

**Table S9:** Empirical power with 46,887 first-degree white British relatives under the homogeneous model with normal error distribution. Power is reported at different significance levels.

| | Quantiles | $10^{-8}$ | $10^{-7}$ | $10^{-6}$ | $10^{-5}$ | 0.0001 | 0.001 | 0.005 | 0.01 | 0.05 |
| --- | --- | --- | --- | --- | --- | --- | --- | --- | --- | --- |
| Regenie | - | 0.0116 | 0.0185 | 0.0299 | 0.0494 | 0.0822 | 0.1405 | 0.2082 | 0.2478 | 0.3779 |
| QRS | composite | 0.0036 | 0.0067 | 0.0128 | 0.0246 | 0.0489 | 0.1000 | 0.1669 | 0.2088 | 0.3524 |
|  | 0.2 | 0.0011 | 0.0023 | 0.0050 | 0.0108 | 0.0241 | 0.0557 | 0.1018 | 0.1330 | 0.2519 |
|  | 0.5 | 0.0028 | 0.0052 | 0.0100 | 0.0194 | 0.0388 | 0.0796 | 0.1337 | 0.1684 | 0.2934 |
|  | 0.8 | 0.0012 | 0.0024 | 0.0050 | 0.0108 | 0.0240 | 0.0555 | 0.1015 | 0.1328 | 0.2517 |
| Regenie.QRS | composite | 0.0037 | 0.0068 | 0.0127 | 0.0246 | 0.0488 | 0.0992 | 0.1662 | 0.2083 | 0.3509 |
|  | 0.2 | 0.0012 | 0.0024 | 0.0050 | 0.0109 | 0.0242 | 0.0555 | 0.1014 | 0.1323 | 0.2516 |
|  | 0.5 | 0.0029 | 0.0054 | 0.0100 | 0.0195 | 0.0388 | 0.0795 | 0.1341 | 0.1687 | 0.2935 |
|  | 0.8 | 0.0012 | 0.0025 | 0.0051 | 0.0110 | 0.0241 | 0.0554 | 0.1014 | 0.1322 | 0.2507 |
| BLUP.QRS | composite | 0.0033 | 0.0062 | 0.0118 | 0.0230 | 0.0464 | 0.0955 | 0.1618 | 0.2032 | 0.3458 |
|  | 0.2 | 0.0010 | 0.0021 | 0.0045 | 0.0099 | 0.0228 | 0.0534 | 0.0982 | 0.1287 | 0.2465 |
|  | 0.5 | 0.0026 | 0.0049 | 0.0094 | 0.0184 | 0.0367 | 0.0761 | 0.1295 | 0.1635 | 0.2886 |
|  | 0.8 | 0.0011 | 0.0022 | 0.0046 | 0.0100 | 0.0227 | 0.0529 | 0.0980 | 0.1286 | 0.2465 |

**Table S10:**
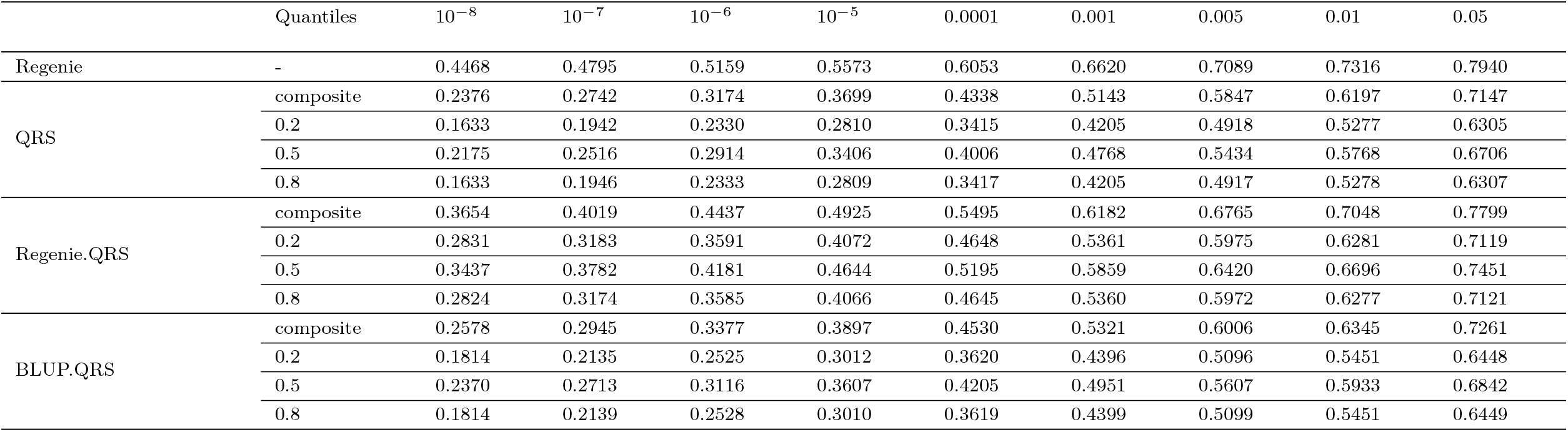
Empirical power with 409,627 white British individuals under the homogenous model with normal error distribution. Power is reported at different significance levels.

**Table S11:** Empirical power with 46,887 first-degree white British relatives under the *t*_(2)_ error model. Power is reported at different significance levels.

| | Quantiles | $10^{-8}$ | $10^{-7}$ | $10^{-6}$ | $10^{-5}$ | 0.0001 | 0.001 | 0.005 | 0.01 | 0.05 |
| --- | --- | --- | --- | --- | --- | --- | --- | --- | --- | --- |
| Regenie | - | 0.0133 | 0.0210 | 0.0331 | 0.0534 | 0.0878 | 0.1473 | 0.2153 | 0.2551 | 0.3853 |
| QRS | composite | 0.0057 | 0.0100 | 0.0178 | 0.0326 | 0.0609 | 0.1166 | 0.1873 | 0.2305 | 0.3742 |
|  | 0.2 | 0.0018 | 0.0033 | 0.0068 | 0.0138 | 0.0295 | 0.0643 | 0.1134 | 0.1463 | 0.2686 |
|  | 0.5 | 0.0047 | 0.0082 | 0.0148 | 0.0271 | 0.0506 | 0.0971 | 0.1559 | 0.1926 | 0.3202 |
|  | 0.8 | 0.0017 | 0.0034 | 0.0067 | 0.0138 | 0.0296 | 0.0646 | 0.1142 | 0.1466 | 0.2684 |
| Regenie.QRS | composite | 0.0077 | 0.0131 | 0.0226 | 0.0394 | 0.0706 | 0.1296 | 0.2019 | 0.2454 | 0.3883 |
|  | 0.2 | 0.0023 | 0.0044 | 0.0083 | 0.0167 | 0.0341 | 0.0716 | 0.1235 | 0.1570 | 0.2799 |
|  | 0.5 | 0.0060 | 0.0102 | 0.0191 | 0.0335 | 0.0600 | 0.1105 | 0.1724 | 0.2100 | 0.3390 |
|  | 0.8 | 0.0023 | 0.0044 | 0.0084 | 0.0166 | 0.0340 | 0.0717 | 0.1236 | 0.1570 | 0.2803 |
| BLUP.QRS | composite | 0.0055 | 0.0097 | 0.0175 | 0.0319 | 0.0596 | 0.1145 | 0.1846 | 0.2271 | 0.3706 |
|  | 0.2 | 0.0017 | 0.0032 | 0.0064 | 0.0134 | 0.0285 | 0.0627 | 0.1112 | 0.1435 | 0.2650 |
|  | 0.5 | 0.0046 | 0.0081 | 0.0145 | 0.0267 | 0.0497 | 0.0959 | 0.1545 | 0.1911 | 0.3185 |
|  | 0.8 | 0.0017 | 0.0032 | 0.0064 | 0.0132 | 0.0283 | 0.0631 | 0.1120 | 0.1444 | 0.2651 |

**Table S12:** Empirical power with 409,627 white British individuals under the *t*_(2)_ error model. Power is reported at different significance levels.

| | Quantiles | $10^{-8}$ | $10^{-7}$ | $10^{-6}$ | $10^{-5}$ | 0.0001 | 0.001 | 0.005 | 0.01 | 0.05 |
| --- | --- | --- | --- | --- | --- | --- | --- | --- | --- | --- |
| Regenie | - | 0.4618 | 0.4940 | 0.5298 | 0.5706 | 0.6174 | 0.6725 | 0.7182 | 0.7402 | 0.8008 |
| QRS | composite | 0.2692 | 0.3057 | 0.3489 | 0.4006 | 0.4630 | 0.5406 | 0.6079 | 0.6411 | 0.7307 |
|  | 0.2 | 0.1841 | 0.2166 | 0.2560 | 0.3047 | 0.3655 | 0.4435 | 0.5134 | 0.5487 | 0.6481 |
|  | 0.5 | 0.2521 | 0.2865 | 0.3271 | 0.3763 | 0.4355 | 0.5095 | 0.5736 | 0.6054 | 0.6942 |
|  | 0.8 | 0.1839 | 0.2162 | 0.2554 | 0.3041 | 0.3649 | 0.4437 | 0.5134 | 0.5488 | 0.6483 |
| Regenie.QRS | composite | 0.4497 | 0.4836 | 0.5216 | 0.5646 | 0.6144 | 0.6740 | 0.7235 | 0.7476 | 0.8117 |
|  | 0.2 | 0.3491 | 0.3841 | 0.4238 | 0.4702 | 0.5245 | 0.5905 | 0.6460 | 0.6732 | 0.7483 |
|  | 0.5 | 0.4356 | 0.4688 | 0.5057 | 0.5480 | 0.5964 | 0.6542 | 0.7025 | 0.7258 | 0.7894 |
|  | 0.8 | 0.3490 | 0.3839 | 0.4238 | 0.4702 | 0.5244 | 0.5904 | 0.6457 | 0.6731 | 0.7482 |
| BLUP.QRS | composite | 0.2921 | 0.3288 | 0.3716 | 0.4220 | 0.4830 | 0.5585 | 0.6233 | 0.6552 | 0.7416 |
|  | 0.2 | 0.2038 | 0.2369 | 0.2766 | 0.3254 | 0.3862 | 0.4633 | 0.5311 | 0.5654 | 0.6620 |
|  | 0.5 | 0.2746 | 0.3095 | 0.3502 | 0.3985 | 0.4568 | 0.5289 | 0.5909 | 0.6215 | 0.7068 |
|  | 0.8 | 0.2037 | 0.2369 | 0.2768 | 0.3255 | 0.3857 | 0.4631 | 0.5312 | 0.5655 | 0.6619 |

**Table S13:**
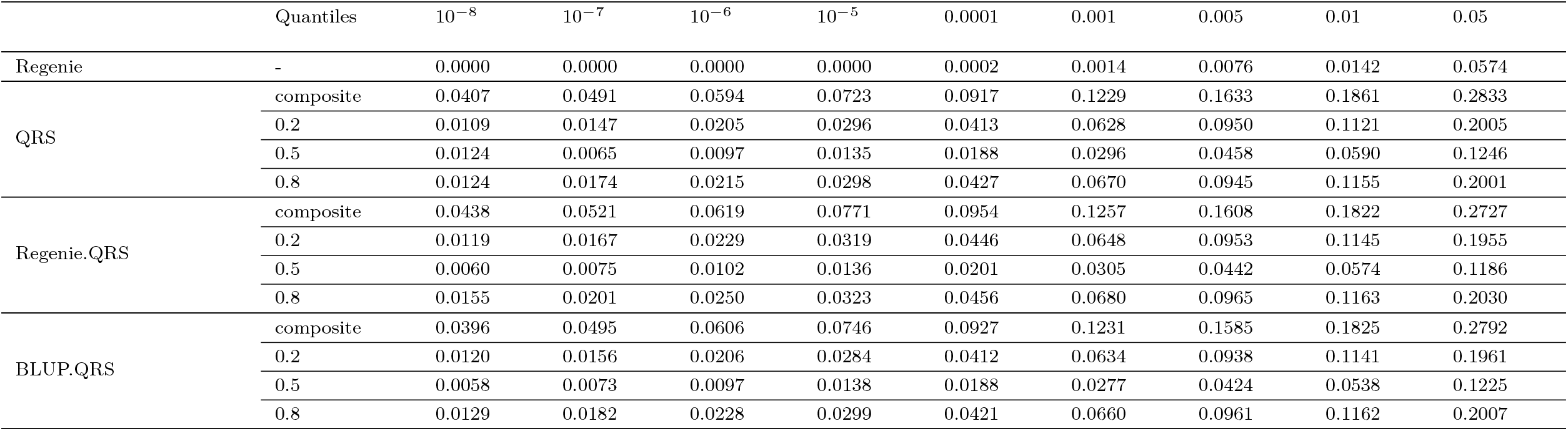
Empirical power with 46,887 first-degree white British relatives under the dominance model. Power is reported at different significance levels.

**Table S14:** Empirical power with 409,627 white British individuals under the dominance model. Power is reported at different significance levels.

| | Quantiles | $10^{-8}$ | $10^{-7}$ | $10^{-6}$ | $10^{-5}$ | 0.0001 | 0.001 | 0.005 | 0.01 | 0.05 |
| --- | --- | --- | --- | --- | --- | --- | --- | --- | --- | --- |
| Regenie | - | 0.0005 | 0.0005 | 0.0008 | 0.0012 | 0.0023 | 0.0054 | 0.0131 | 0.0215 | 0.0711 |
| QRS | composite | 0.2000 | 0.2167 | 0.2353 | 0.2558 | 0.2838 | 0.3217 | 0.3654 | 0.3915 | 0.4815 |
|  | 0.2 | 0.1276 | 0.1419 | 0.1575 | 0.1794 | 0.2100 | 0.2509 | 0.2918 | 0.3166 | 0.4042 |
|  | 0.5 | 0.0520 | 0.0578 | 0.0644 | 0.0724 | 0.0843 | 0.1042 | 0.1264 | 0.1391 | 0.2067 |
|  | 0.8 | 0.1272 | 0.1411 | 0.1588 | 0.1814 | 0.2091 | 0.2545 | 0.2972 | 0.3218 | 0.4076 |
| Regenie.QRS | composite | 0.2504 | 0.2673 | 0.2853 | 0.3111 | 0.3434 | 0.3816 | 0.4267 | 0.4492 | 0.5321 |
|  | 0.2 | 0.1698 | 0.1858 | 0.2064 | 0.2313 | 0.2657 | 0.3080 | 0.3521 | 0.3755 | 0.4610 |
|  | 0.5 | 0.0773 | 0.0842 | 0.0937 | 0.1034 | 0.1181 | 0.1398 | 0.1623 | 0.1771 | 0.2432 |
|  | 0.8 | 0.1650 | 0.1829 | 0.2042 | 0.2299 | 0.2642 | 0.3083 | 0.3495 | 0.3721 | 0.4522 |
| BLUP.QRS | composite | 0.2101 | 0.2264 | 0.2431 | 0.2669 | 0.2967 | 0.3359 | 0.3768 | 0.4032 | 0.4863 |
|  | 0.2 | 0.1358 | 0.1505 | 0.1697 | 0.1910 | 0.2220 | 0.2606 | 0.3061 | 0.3295 | 0.4123 |
|  | 0.5 | 0.0563 | 0.0618 | 0.0693 | 0.0774 | 0.0891 | 0.1112 | 0.1334 | 0.1469 | 0.2131 |
|  | 0.8 | 0.1336 | 0.1476 | 0.1671 | 0.1911 | 0.2208 | 0.2629 | 0.3074 | 0.3322 | 0.4157 |

**Table S15:**
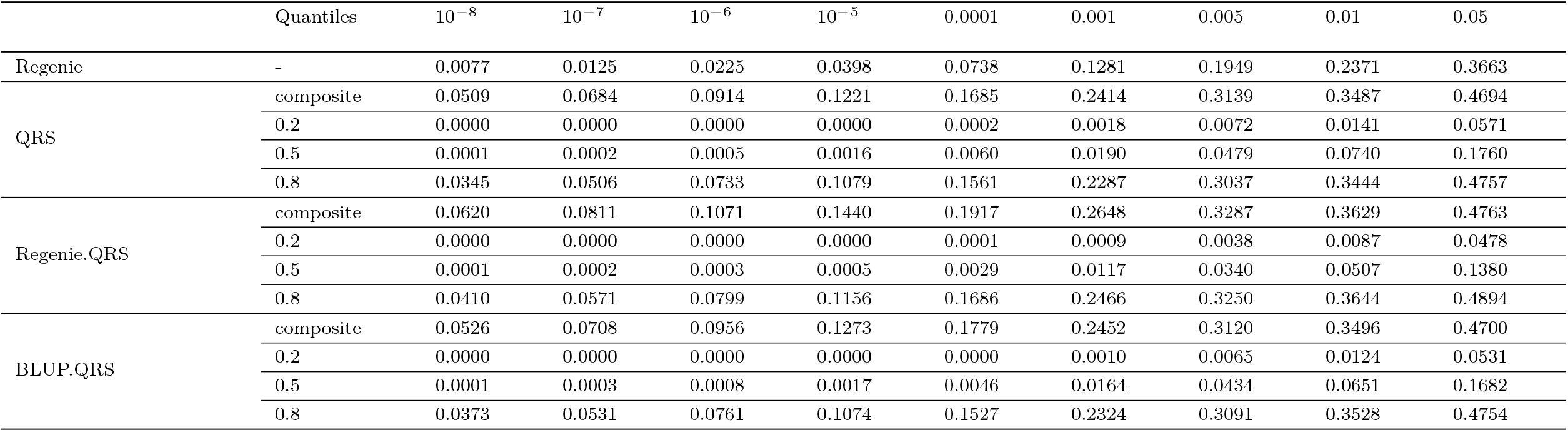
Empirical power with 46,887 first-degree white British relatives under the heterogeneous model. Power is reported at different significance levels.

**Table S16:** Empirical power with 409,627 white British individuals under the heterogenous model. Power is reported at different significance levels.

| | Quantiles | $10^{-8}$ | $10^{-7}$ | $10^{-6}$ | $10^{-5}$ | 0.0001 | 0.001 | 0.005 | 0.01 | 0.05 |
| --- | --- | --- | --- | --- | --- | --- | --- | --- | --- | --- |
| Regenie | - | 0.3849 | 0.421 | 0.4593 | 0.507 | 0.5558 | 0.6164 | 0.6684 | 0.6942 | 0.7612 |
| QRS | composite | 0.4859 | 0.5154 | 0.5446 | 0.579 | 0.6194 | 0.6661 | 0.7073 | 0.7241 | 0.7788 |
|  | 0.2 | 0 | 0 | 0.0001 | 0.0002 | 0.0004 | 0.0038 | 0.0125 | 0.0205 | 0.0798 |
|  | 0.5 | 0.0727 | 0.0944 | 0.1229 | 0.1587 | 0.2125 | 0.2879 | 0.3654 | 0.4063 | 0.5242 |
|  | 0.8 | 0.4453 | 0.4758 | 0.5134 | 0.5523 | 0.5994 | 0.653 | 0.6985 | 0.7193 | 0.7847 |
| Regenie.QRS | composite | 0.58 | 0.601 | 0.6271 | 0.6571 | 0.6902 | 0.7297 | 0.7613 | 0.7777 | 0.8218 |
|  | 0.2 | 0 | 0 | 0 | 0 | 0.0002 | 0.0009 | 0.0067 | 0.0108 | 0.0528 |
|  | 0.5 | 0.0274 | 0.0403 | 0.0568 | 0.0851 | 0.1251 | 0.1898 | 0.26 | 0.2982 | 0.4273 |
|  | 0.8 | 0.5389 | 0.5661 | 0.5965 | 0.6307 | 0.6708 | 0.7154 | 0.755 | 0.775 | 0.8303 |
| BLUP.QRS | composite | 0.5041 | 0.5291 | 0.5602 | 0.5941 | 0.6312 | 0.6793 | 0.7144 | 0.7355 | 0.7876 |
|  | 0.2 | 0 | 0 | 0.0001 | 0.0001 | 0.0001 | 0.0028 | 0.0096 | 0.0165 | 0.0703 |
|  | 0.5 | 0.0661 | 0.0869 | 0.1156 | 0.1533 | 0.2053 | 0.2815 | 0.3531 | 0.3956 | 0.5181 |
|  | 0.8 | 0.4601 | 0.4914 | 0.5283 | 0.5708 | 0.6119 | 0.6613 | 0.7093 | 0.7325 | 0.793 |

**Table S17:** Empirical type I error rates with 10,000 first-degree white British relatives under the homogenous model.

| | Quantiles | $10^{-8}$ | $10^{-7}$ | $10^{-6}$ | $10^{-5}$ | $10^{-4}$ | 0.001 | 0.005 | 0.01 | 0.05 |
| --- | --- | --- | --- | --- | --- | --- | --- | --- | --- | --- |
| | | $(0, 5.74 \times 10^{-8})$ | $(0, 2.50 \times 10^{-7})$ | $(5.26 \times 10^{-7}, 1.47 \times 10^{-6})$ | $(8.50 \times 10^{-6}, 1.15 \times 10^{-5})$ | $(9.53 \times 10^{-5}, 1.05 \times 10^{-4})$ | $(0.0000985, 0.00101)$ | $(0.00497, 0.00503)$ | $(0.00995, 0.01005)$ | $(0.0499, 0.0501)$ |
| Regenie | - | $< 5.84 \times 10^{-8}$ | $< 5.84 \times 10^{-8}$ | $4.09 \times 10^{-7}$ | $6.95 \times 10^{-6}$ | $5.99 \times 10^{-5}$ | 0.0007 | 0.0036 | 0.0076 | 0.0423 |
| QRS | composite | $1.17 \times 10^{-7}$ | $1.11 \times 10^{-6}$ | $5.96 \times 10^{-6}$ | $4.22 \times 10^{-5}$ | $3.27 \times 10^{-4}$ | 0.0025 | 0.0107 | 0.0198 | 0.0808 |
| | 0.2 | $< 5.84 \times 10^{-8}$ | $1.17 \times 10^{-7}$ | $3.27 \times 10^{-6}$ | $2.99 \times 10^{-5}$ | $2.39 \times 10^{-4}$ | 0.0019 | 0.0081 | 0.0151 | 0.0644 |
| | 0.5 | $1.17 \times 10^{-7}$ | $1.05 \times 10^{-6}$ | $4.97 \times 10^{-6}$ | $3.89 \times 10^{-5}$ | $2.80 \times 10^{-4}$ | 0.0021 | 0.0087 | 0.0160 | 0.0668 |
| | 0.8 | $1.75 \times 10^{-7}$ | $4.67 \times 10^{-7}$ | $3.80 \times 10^{-6}$ | $3.03 \times 10^{-5}$ | $2.43 \times 10^{-4}$ | 0.0019 | 0.0081 | 0.0151 | 0.0643 |
| Regenie.QRS | composite | $< 5.84 \times 10^{-8}$ | $< 5.84 \times 10^{-8}$ | $5.26 \times 10^{-7}$ | $7.83 \times 10^{-6}$ | $7.90 \times 10^{-5}$ | 0.0008 | 0.0045 | 0.0093 | 0.0493 |
| | 0.2 | $< 5.84 \times 10^{-8}$ | $< 5.84 \times 10^{-8}$ | $7.60 \times 10^{-7}$ | $7.95 \times 10^{-6}$ | $7.78 \times 10^{-5}$ | 0.0008 | 0.0043 | 0.0089 | 0.0465 |
| | 0.5 | $< 5.84 \times 10^{-8}$ | $1.17 \times 10^{-7}$ | $7.01 \times 10^{-7}$ | $5.43 \times 10^{-6}$ | $6.08 \times 10^{-5}$ | 0.0007 | 0.0038 | 0.0080 | 0.0436 |
| | 0.8 | $< 5.84 \times 10^{-8}$ | $5.84 \times 10^{-8}$ | $4.09 \times 10^{-7}$ | $7.60 \times 10^{-6}$ | $8.06 \times 10^{-5}$ | 0.0008 | 0.0043 | 0.0089 | 0.0465 |
| BLUP.QRS | composite | $< 5.84 \times 10^{-8}$ | $5.84 \times 10^{-8}$ | $1.05 \times 10^{-6}$ | $1.25 \times 10^{-5}$ | $1.22 \times 10^{-4}$ | 0.0012 | 0.0060 | 0.0119 | 0.0583 |
| | 0.2 | $< 5.84 \times 10^{-8}$ | $5.84 \times 10^{-8}$ | $9.35 \times 10^{-7}$ | $1.03 \times 10^{-5}$ | $1.07 \times 10^{-4}$ | 0.0011 | 0.0053 | 0.0104 | 0.0513 |
| | 0.5 | $< 5.84 \times 10^{-8}$ | $1.17 \times 10^{-7}$ | $1.17 \times 10^{-6}$ | $1.16 \times 10^{-5}$ | $1.10 \times 10^{-4}$ | 0.0011 | 0.0053 | 0.0106 | 0.0517 |
| | 0.8 | $< 5.84 \times 10^{-8}$ | $1.75 \times 10^{-7}$ | $1.58 \times 10^{-6}$ | $1.23 \times 10^{-5}$ | $1.12 \times 10^{-4}$ | 0.0011 | 0.0052 | 0.0104 | 0.0512 |

**Table S18:** Empirical power with 10,000 first-degree white British relatives under the homogenous model. Power is reported at different significance levels.

| | Quantiles | $10^{-8}$ | $10^{-7}$ | $10^{-6}$ | $10^{-5}$ | 0.0001 | 0.001 | 0.005 | 0.01 | 0.05 |
| --- | --- | --- | --- | --- | --- | --- | --- | --- | --- | --- |
| Regenie | - | $1.80 \times 10^{-5}$ | $5.70 \times 10^{-5}$ | $1.75 \times 10^{-4}$ | 0.0006 | 0.0025 | 0.0101 | 0.0278 | 0.0433 | 0.1229 |
| QRS | composite | $2.40 \times 10^{-5}$ | $6.10 \times 10^{-5}$ | $2.18 \times 10^{-4}$ | 0.0008 | 0.0033 | 0.0139 | 0.0381 | 0.0592 | 0.1605 |
| | 0.2 | $1.00 \times 10^{-6}$ | $1.60 \times 10^{-5}$ | $8.80 \times 10^{-5}$ | 0.0004 | 0.0017 | 0.0077 | 0.0228 | 0.0366 | 0.1110 |
| | 0.5 | $2.10 \times 10^{-5}$ | $5.70 \times 10^{-5}$ | $1.85 \times 10^{-4}$ | 0.0007 | 0.0027 | 0.0107 | 0.0291 | 0.0453 | 0.1270 |
| | 0.8 | $9.00 \times 10^{-6}$ | $2.40 \times 10^{-5}$ | $8.20 \times 10^{-5}$ | 0.0004 | 0.0016 | 0.0077 | 0.0228 | 0.0367 | 0.1118 |
| regenie.QRS | composite | $1.00 \times 10^{-6}$ | $9.00 \times 10^{-6}$ | $4.50 \times 10^{-5}$ | 0.0002 | 0.0012 | 0.0065 | 0.0215 | 0.0361 | 0.1171 |
| | 0.2 | $1.00 \times 10^{-6}$ | $4.00 \times 10^{-6}$ | $2.10 \times 10^{-5}$ | 0.0001 | 0.0007 | 0.0041 | 0.0140 | 0.0241 | 0.0855 |
| | 0.5 | $1.00 \times 10^{-6}$ | $7.00 \times 10^{-6}$ | $4.70 \times 10^{-5}$ | 0.0002 | 0.0010 | 0.0052 | 0.0167 | 0.0282 | 0.0941 |
| | 0.8 | $1.00 \times 10^{-6}$ | $6.00 \times 10^{-6}$ | $2.20 \times 10^{-5}$ | 0.0001 | 0.0007 | 0.0040 | 0.0141 | 0.0242 | 0.0856 |
| BLUP.QRS | composite | $5.00 \times 10^{-6}$ | $2.30 \times 10^{-5}$ | $8.10 \times 10^{-5}$ | 0.0003 | 0.0016 | 0.0080 | 0.0256 | 0.0418 | 0.1293 |
| | 0.2 | $3.00 \times 10^{-6}$ | $5.00 \times 10^{-6}$ | $3.50 \times 10^{-5}$ | 0.0002 | 0.0009 | 0.0048 | 0.0160 | 0.0269 | 0.0923 |
| | 0.5 | $1.00 \times 10^{-6}$ | $1.90 \times 10^{-5}$ | $7.10 \times 10^{-5}$ | 0.0003 | 0.0014 | 0.0066 | 0.0205 | 0.0332 | 0.1046 |
| | 0.8 | $1.00 \times 10^{-6}$ | $6.00 \times 10^{-6}$ | $2.60 \times 10^{-5}$ | 0.0002 | 0.0009 | 0.0048 | 0.0160 | 0.0269 | 0.0922 |

**Table S19:** Comparison of power between QRS based on 325,667 unrelated individuals and Regenie.QRS based on 409,627 individuals including related samples. Power is reported at different significance levels, under homogeneous model.

| | Quantiles | $10^{-8}$ | $10^{-7}$ | $10^{-6}$ | $10^{-5}$ | 0.0001 | 0.001 | 0.005 | 0.01 | 0.05 |
| --- | --- | --- | --- | --- | --- | --- | --- | --- | --- | --- |
| QRS | composite | 0.1878 | 0.2224 | 0.2647 | 0.3165 | 0.3825 | 0.4667 | 0.5418 | 0.5796 | 0.6831 |
|  | 0.2 | 0.1213 | 0.1489 | 0.1847 | 0.2301 | 0.2901 | 0.3701 | 0.4442 | 0.4826 | 0.5930 |
|  | 0.5 | 0.1694 | 0.2012 | 0.2399 | 0.2883 | 0.3490 | 0.4275 | 0.4978 | 0.5336 | 0.6361 |
|  | 0.8 | 0.1213 | 0.1493 | 0.1846 | 0.2303 | 0.2900 | 0.3701 | 0.4445 | 0.4826 | 0.5931 |
| Regenie.QRS | composite | 0.3654 | 0.4019 | 0.4437 | 0.4925 | 0.5495 | 0.6182 | 0.6765 | 0.7048 | 0.7799 |
|  | 0.2 | 0.2831 | 0.3183 | 0.3591 | 0.4071 | 0.4648 | 0.5361 | 0.5975 | 0.6281 | 0.7119 |
|  | 0.5 | 0.3437 | 0.3782 | 0.4181 | 0.4644 | 0.5195 | 0.5859 | 0.6420 | 0.6696 | 0.7451 |
|  | 0.8 | 0.2824 | 0.3174 | 0.3585 | 0.4066 | 0.4645 | 0.5360 | 0.5972 | 0.6277 | 0.7121 |

**Table S20:** Computational time and memory usage across Regenie, QRS, Regenie.QRS, and BLUP.QRS. *N* : sample size; *M*_*b*_: number of SNPs in a genotype block in Regenie; *B*: total number of genotype blocks; *M* : number of variants tested in step 2; *J*_0_: number of ridge parameters at level 0 (step 1 in Regenie); *P* : number of phenotypes; *J*_1_: number of ridge parameters at level 1 (step 1 in Regenie); *R* = *J*_0_*B*: number of predictors from level 0 models; *Q*: number of quantile levels; *S*: number of SNPs used to compute the GRM.

| Method | Time Complexity |  | BMI in UK Biobank |  |
| --- | --- | --- | --- | --- |
|  | Step 1 | Step 2 | Time (CPU hrs) | Memory (GB) |
| Regenie | $O(BNM_b^2) + O(BM_b^3 J_0)$<br>$+O(BNM_b P) + O(NR^2 P)$<br>$+O(R^3 J_1 P)$ | $O(NMP)$ | 41 | 9 |
| QRS | $O(NMQP)$ | | 1144 | 77 |
| Regenie.QRS | $O(BNM_b^2) + O(BM_b^3 J_0)$<br>$+O(BNM_b P) + O(NR^2 P)$<br>$+O(R^3 J_1 P)$ | $O(NMQP)$ | 660 | 76 |
| BLUP.QRS | $O(SN^{1.5})$ | $O(NMQP)$ | 599 | 77 |

